# Unique molecular identifiers don’t need to be unique: a collision-aware estimator for RNA-seq quantification

**DOI:** 10.1101/2025.09.08.674884

**Authors:** Dylan Agyemang, Rafael A. Irizarry, Tavor Z. Baharav

## Abstract

RNA-sequencing (RNA-seq) relies on Unique Molecular Identifiers (UMIs) to accurately quantify gene expression after PCR amplification. Longer UMIs minimize collisions—where two distinct transcripts are assigned the same UMI—at the expense of increased sequencing and synthesis costs. However, it is not clear how long UMIs need to be in practice, especially given the nonuniformity of the empirical UMI distribution. In this work, we develop a method-of-moments estimator that accounts for UMI collisions, accurately quantifying gene expression and preserving downstream biological insights. We show that UMIs need not be unique: shorter UMIs can be used with a more sophisticated estimator.

## 1 Introduction

RNA-sequencing is widely used to study the sequence and abundance of mRNA molecules within cells [1, 2]. In order to generate sufficient genomic material for sequencing, especially at single-cell resolution [3], mRNA molecules must be PCR amplified. However, the nonuniform amplification of PCR makes UMIs essential for accurate quantification [4]. UMIs are randomly generated nucleotide sequences of a specific length that are attached to the reverse-transcribed cDNA molecules in the sequencing process [5]. After PCR amplification, reads that originate from the same molecule will carry the same UMI (Figure 1a). Genomic analysis pipelines [3] identify reads with the same sequence and same UMI as duplicates, and collapse them into a single count. Reads with distinct UMIs but identical sequences are counted separately (Figure 1b). However, the reliability of UMIs to correctly make this distinction depends on the length of the UMI used. At each position of a UMI, we have a choice of 4 nucleotides, yielding a total of *K* = 4^*k*^ distinct length-*k* UMIs. With shorter UMI lengths we observe *collisions*, where multiple identical mRNA transcripts are assigned the same UMI sequence [6] (Figure 1c). Consequently, deduplication leads to an underestimation of the true transcript abundance, as it misattributes the collision to a PCR duplicate rather than a distinct additional transcript (Figure 1d). With longer UMIs we avoid these collisions, at the expense of additional sequencing cycles and increased synthesis difficulty. As it stands, there is no consensus on how to choose UMI length to balance cost and quantification accuracy, so varied lengths are used in practice. For example, the UMI length was notably increased from 10-bp in 10x Chromium v2 to 12-bp in 10x Chromium v3, while 8-bp was originally used in scRNA-seq [7] and 6-bp or 8-bp is often still used in miRNA experiments [8].

**Fig. 1:**
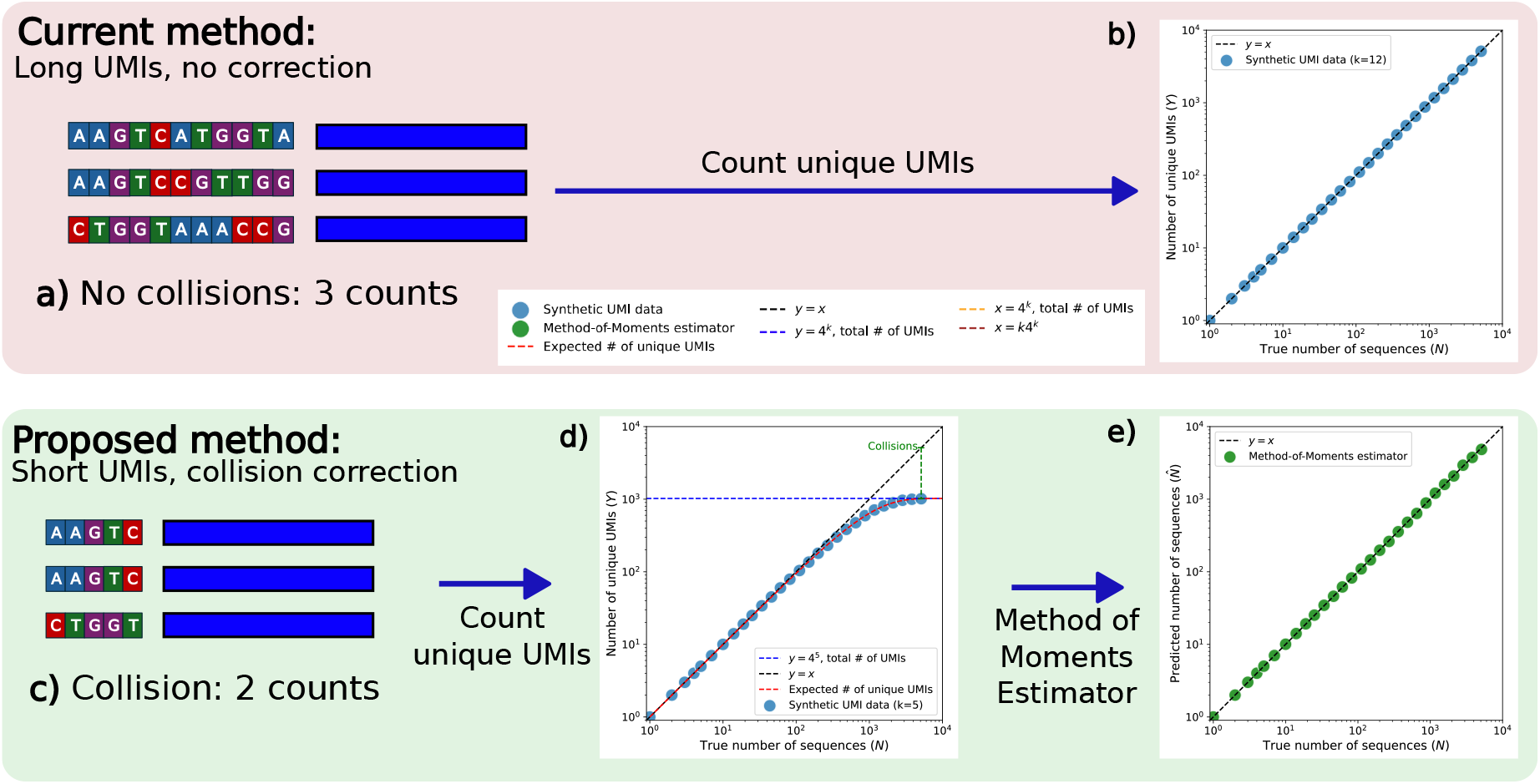
Comparison between the standard pipeline and our collision-aware estimator. Synthetic data, details in Section S1.6. Collisions occur when distinct transcripts are assigned identical UMI sequences. **a)** With long UMIs (*k* = 12) collisions are rare. All 3 transcripts (dark blue rectangles) are assigned different UMIs, and so deduplication yields 3 counts. **b)** Deduplicated UMI counts accurately predict the true number of transcripts for long UMIs. **c)** With short UMIs (*k* = 5) collisions will occur. Here, 3 transcripts are sequenced, but 2 are assigned the same UMI, leading to a deduplicated count of 2. **d)** Deduplicated UMI counts fall far below the *y* = *x* line due to collisions. Our statistical model (red line) accurately predicts this observed relationship. **e)** Our method-of-moments estimator applied to **d** corrects for these collisions.

Several prior works have studied the statistical and computational challenges arising from UMIs. The classical occupancy framework underlying UMI collisions was introduced by Fu et al. [6], who derived the expected number of unique labels (UMIs) observed as a function of the number of molecules and the label pool size, assuming a uniform distribution over labels. Kivioja et al. [5] formalized UMIs as a general tool for absolute molecular quantification in genome-scale sequencing, demonstrating their utility in digital karyotyping and mRNA-seq. Clement et al. [9] formalized collision probabilities and allelic fraction distortion for amplicon sequencing, providing guidelines for choosing UMI lengths that avoid collisions entirely, but did not develop an estimator that corrects for collisions after the fact. Substantial effort has been devoted to the upstream problem of UMI error correction: graph-based approaches [10], alignment-free methods [11], Phred quality-based methods [12], and Hamming distance-based deduplication [5, 7]. However, error correction methods are not designed to account for collisions, and can even exacerbate underestimation with short UMI lengths by over-collapsing UMIs that are close in Hamming distance but actually represent distinct transcripts (Section S1.3). The most comprehensive prior treatment is dropEst [13], which jointly models UMI error correction, deduplication, and collision adjustment under a nonuniform UMI distribution. However, the dropEst model is computationally intensive, requiring quantization and approximate computation. It does not admit a rigorous theoretical analysis of estimator performance or estimation limits, and because it operates as an end-to-end pipeline from raw reads to final count matrix, it cannot be applied as a modular plug-in to standard pipelines or benchmarked on synthetically shortened UMIs. Recent work has also identified that the empirical nonuniformity of UMI distributions arises in part from truncated oligonucleotide synthesis on beads [14], a mechanistic insight we incorporate into our statistical model (Section 4.1).

In this work, we address these gaps and develop a method-of-moments estimator for the true transcript abundance that accounts for UMI collisions, is near-optimal, is computationally efficient, and is designed as a modular plug-in for existing pipelines. Expanding on the basic UMI collision model [6], we derive a simple closed-form estimator in the uniform case and an efficiently computable estimator in the nonuniform case, requiring only a one-time lookup table precomputation. Leveraging classical statistical results [15], we show that our estimator essentially matches the Cramér–Rao lower bound in a simplified binomial setting (Theorem 1). In the Poissonized setting, we derive the maximum likelihood estimator (MLE), which uses information about *which* UMIs were observed, not just the number of unique ones, and show that our method-of-moments estimator is optimal when ***p*** is uniform, with a characterizable efficiency gap that grows with the skewness of ***p*** (Section S3). However, in practice the method-of-moments estimator performs essentially as well as the MLE, in a much more computationally efficient manner (Sections S3 and S4). We characterize the threshold for nontrivial estimation, showing that beyond the coupon collector [16] threshold of *K* log *K* transcripts, accurate estimation is impossible (Proposition 2). We provide asymptotic confidence intervals via the delta method (Section S5.2). Going beyond the uniform assumption of prior works, we develop a mechanistic statistical model of UMI nonuniformity that accounts for failures in UMI synthesis (Section 4.1). Our estimator is computationally efficient (precompute lookup table, then constant time evaluation) and works in concert with existing UMI error correction methods, where error correction serves as a preprocessing step and our collision adjustment is applied afterward. This modularity, absent from prior collision-aware approaches, enables straightforward integration into any existing RNA-seq pipeline and allows us to validate performance on synthetically shortened UMIs. Our estimator allows researchers to reclaim 6 base pairs from the UMI for biological payload or additional barcodes, without losing any quantification accuracy. This can be critical in spatial barcoding and combinatorial indexing, which require extensive sequencing real estate.

## 2 Problem formulation

Methodologically, in this work we study, for a given gene, how to estimate the number of unique pre-amplification mRNA molecules. Specifying to one particular mRNA molecule, we make the assumption that the UMI sequence does not impact its amplification. This assumption is reasonable, as the UMI is a short sequence (up to 12 bp) attached to a much longer cDNA transcript (several hundred base pairs), and is therefore unlikely to impact amplification. This motivates our statistical model where each mRNA molecule is tagged with a UMI randomly drawn from some distribution over the 4^*k*^ possible UMIs, and that each pre-amplification mRNA molecule has an equal probability of appearing in the set of sequenced reads.

Under these assumptions, we model our problem on a gene-by-gene basis using the classical balls and bins framework. Consider *N* identical balls randomly assigned into *K* bins, with the observation *Y* denoting the number of bins with at least one ball. In our sequencing model, this translates to *N* mRNA transcripts for a given gene before PCR amplification, *K* = 4^*k*^ possible UMIs, and *Y* observed unique UMIs. Given the UMI distribution *p* over the *K* UMIs, we want to estimate the number of transcripts *N* after observing *Y* = *y* unique UMIs. This estimation question relates to the classical occupancy problem [15]. Naively, one would estimate 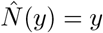, which is suboptimal as for 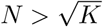 (the birthday paradox [16]) collisions occur with probability greater than 1/2. More importantly, estimation is possible up to the coupon collector threshold of *N* = *K* log *K*, the expected number of original mRNA transcripts that need to be sequenced in order for us to observe all *K* UMIs [16]. We show that *N* cannot be consistently estimated above this threshold, and conversely that below this threshold our estimator provides excellent performance, theoretically matching the Cramér–Rao lower bound in a simplified binomial setting (Section S7). These results simplify nicely when all UMIs are equally likely, but in practice UMIs do not follow a uniform distribution, which biases the estimator if unaccounted for. We adapt our estimator to this nonuniform case, yielding an efficiently computable estimator 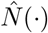 which we derive below.

## 3 Abundance estimator

Recall our statistical model, restated here for completeness: *N* identical balls are randomly assigned into *K* bins, with the observation *Y* denoting the number of bins with at least one ball. In our sequencing model, this translates to *N* mRNA transcripts before PCR amplification, *K* = 4^*k*^ possible UMIs, and *Y* unique UMIs observed. The *N* balls (transcripts) which we denote by *X*_*i*_, follow some distribution *p* over the set [*K*] = {1, 2, …, *K*}, and are independent and identically distributed. Then:

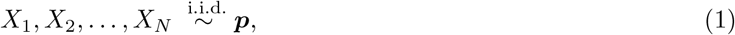

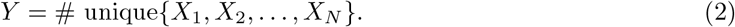

Mathematically, *Y* = |{*X*_1_, *X*_2_, …, *X*_*N*_ }|, where |*S*| denotes the cardinality of a set *S*, in our case the number of distinct UMIs observed. Given *p* and *Y*, we want to estimate *N* . By representing *Y* as a sum of *K* indicators, we can compute its mean and variance.

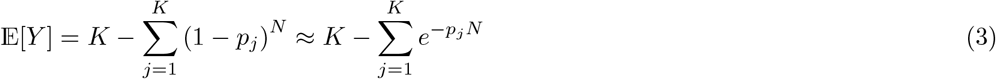

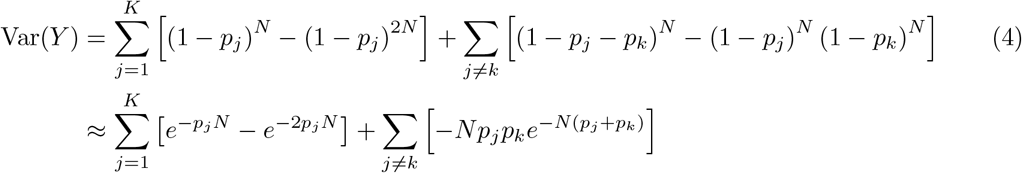

A second order Taylor approximation is necessary to simplify the crossterms in the variance. Observe that 𝔼 [*Y* ] is a concave function of ***p*** for *N* ≥ 1, as can be verified by taking the second derivative. By Jensen’s inequality, this implies that the expected number of unique UMIs observed is maximized when ***p*** is uniform.

This assumption that the *X*_*i*_ are independent and identically distributed need not perfectly hold in practice. For example, sequencing errors may introduce dependencies between the *X*_*i*_ (Section S1.3). Additionally, the UMI distribution may vary across genes, due to binding affinities; however, as we show in Figure 4a, all genes appear to follow the same UMI distribution, except for MALAT1. Overall, this simple model in Equations (1) and (2) fits the data quite well with a common UMI distribution ***p*** in practice, and we discuss estimating ***p*** and identifying the deviation of MALAT1 in Section 4.

### 3.1 Simplification in uniform setting

Simplifying Equation (3) in the case when all UMIs are equally likely we recover a classical result [6]. Rearranging the expectation of *Y*, we obtain our method-of-moments estimator in the uniform UMI setting, when we observe *Y* = *y* unique UMIs:

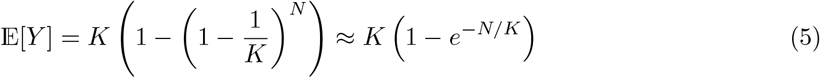

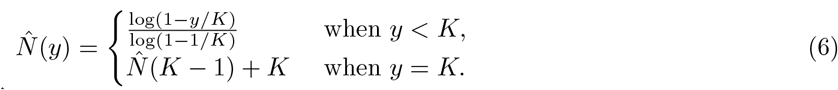

This choice of 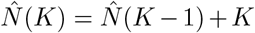 is motivated by the fact that the expected number of additional transcripts required to go from observing *K* − 1 unique UMIs to *K* unique UMIs is *K*. This extension of 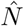 retains the convexity of the estimator (Proposition 1). Extrapolation is an edge case that should be rarely used in practice: if *Y* = *K* is observed then *k* is likely too small for the given sequencing depth.

This balls and bins setting is known to have several interesting thresholds, considering the case of large *K* [16]. First is the classical birthday paradox, which occurs at 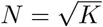. At this point, collisions occur with probability greater than 1/2, indicating that the naive estimator will underestimate the true counts over half the time. Next is the value of *N* = *K*. This is the maximum counts that the naive estimator will output, where we note that at this value of *N*, 𝔼[*Y* ] ≈ (1 − *e*^−1^)*N* ≈ .63*N*, indicating the dramatic bias of the collision-oblivious (naive) estimator. Finally, we have the coupon collector threshold of *N* = *K* log *K*, the expected number of unique mRNA transcripts that need to be sequenced in order for our observations to saturate with *Y* = *K*, where *all K* UMIs are observed. We show that this is the true threshold for consistent estimation, where for *N > K* log *K* we have *Y* = *K* with high probability (Proposition 2). This implies that all *K* UMIs are observed, beyond which larger *N* cannot be estimated. Surprisingly, we show that for *N* ≤ *cK* log *K* for *c <* 1, our estimator performs near optimally, with mean squared error (MSE) matching the Cramér–Rao lower bound [17] in a simplified binomial setting (Theorem 1).

### 3.2 Method-of-moments estimator (nonuniform)

While our estimator in the previous subsection is clean, it does not account for the nonuniformity of UMI distributions in practice (discussed in more detail in Section 4). In the nonuniform case with probabilities 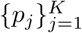 the expected number of unique UMIs of length *k* observed, given that there are

#### Algorithm 1

Collision-aware estimator

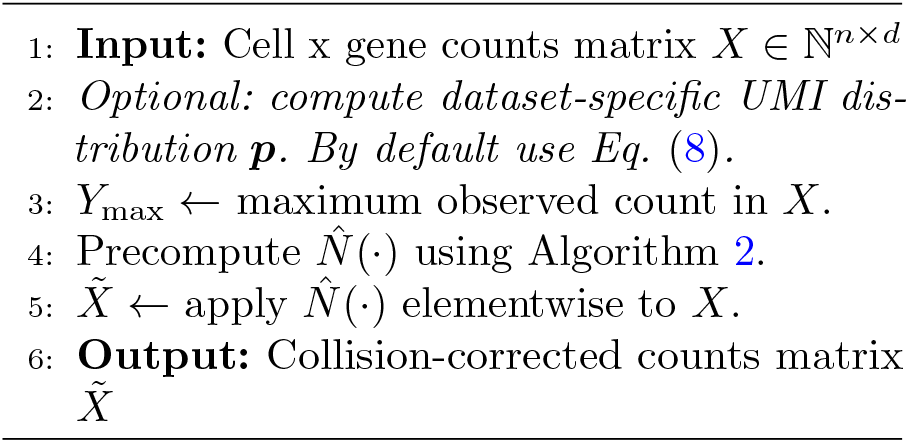

#### Algorithm 2

Precomputation of 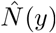

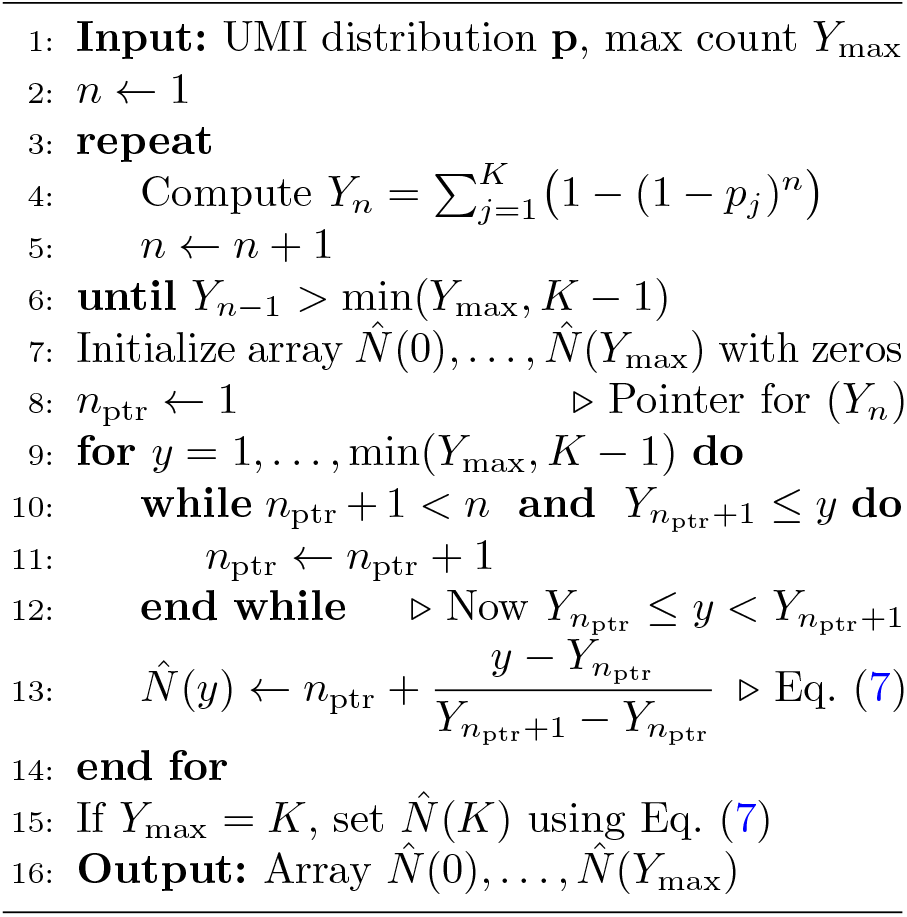

Computation and evaluation of our proposed collision-aware estimator. Algorithm 1 details the overall procedure, while Algorithm 2 describes the precomputation of the estimator 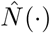.Note that after precomputation, each count can be mapped to its collision-corrected estimate in constant time. Algorithm 1 is applied after UMI error correction and deduplication, before downstream analysis (additional runtime and computational complexity analysis in Section S5.3).

*N* transcripts before PCR amplification, is 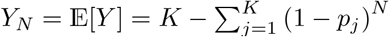 (from Equation (3)). Inverting this expression to solve for *N* as a function of *Y* does not yield a closed form solution. However, given that *Y*_*N*_ is monotonically increasing in *N* (and concave), we can define 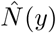 by identifying *n* such that *Y*_*n*_ ≤ *y* ≤ *Y*_*n*+1_, and linearly interpolating. Since *p*_*j*_ *>* 0 for all *j, Y*_*N*_ is concave, strictly increasing in *N*, and twice differentiable. Thus, 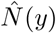 is convex in *y* for *y < K*, as it is the linear interpolation between sampled points of a convex function. As before, the case of *y* = *K* is a priori undefined, as *Y*_*n*_ *< K* for all *n*. Here we use a quadratic extrapolation based on finite differences to yield a simple estimate that retains the convexity of our estimator (details in Section S5.1.1). As before, this extrapolation is an edge case that should be rarely used in practice. Concretely, our final estimator is given by:

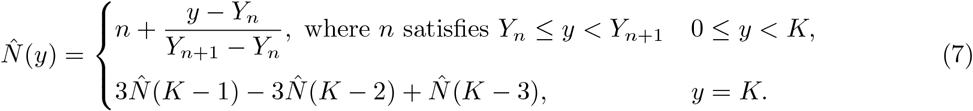

We can approximate the variance of our estimator 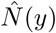 using the delta method. Leveraging the fact that our estimator is (up to linear interpolation) the inverse function of *Y*_*N*_, we obtain that 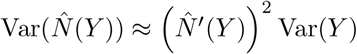 Var(*Y* ). This enables us to compute confidence intervals for our predictions 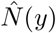 (details in Section S5.2). From classical literature, we know that *Y* is asymptotically normally distributed, as long as Var(*Y* ) → ∞ [15]. This holds for 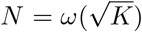, once collisions become non-negligible, up until *N* = *o*(*K* log *K*), before saturation occurs at the coupon collector threshold (details in Section S7). This enables us to quantify the bias of our estimator, which is vanishing for *N* = *o*(*K*) (see Table 1 for full *N* results).

**Table 1:**
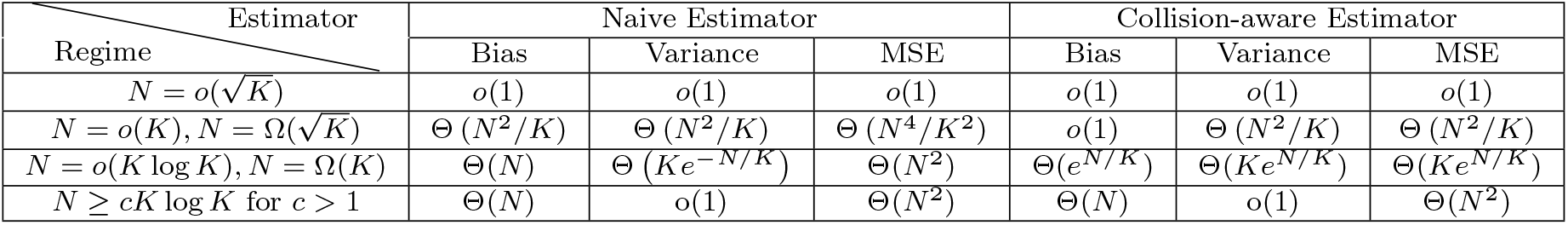
Comparison of bias, variance, and MSE for naive and collision-aware estimators across different regimes of *N* . Uniform UMI distribution assumed for simplicity. Detailed derivations in Section S7.1.

Our estimator can be implemented very efficiently as a modular plug-in to existing RNA-seq pipelines. Given a cell x gene counts matrix *X* ∈ℕ^*n×d*^ of per-gene error-corrected and deduplicated UMI counts, we can compute with Algorithm 1 a collision-corrected counts matrix 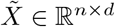. This is done by precomputing the estimator 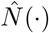 for all observed counts up to the maximum observed count *Y*_max_ in *X* using Algorithm 2, and then applying 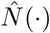 elementwise to *X*. This precomputation step only needs to be done once per dataset, and each count can then be mapped to its collision-corrected estimate in constant time.

## 4 Nonuniformity of empirical UMI distribution

As identified by previous works, UMIs in practice follow a nonuniform distribution [9, 13, 14], which holds consistently across different runs of the same chemistry and so can be estimated from data. This nonuniformity holds across chemistry choices, where drop-Seq UMIs exhibit dramatically higher G frequencies [14]. In this work, we focus primarily on 10X Genomics kits, where this distribution is T-biased (elevated T frequency), and is relatively well approximated by a marginal per-base model, where the probability of the UMI depends only on its aggregate nucleotide composition (Figure S1). The only exception is UMIs with many trailing Ts, which are significantly more likely (Figure 2b,c). Recent work has shown that this is due to *truncated UMI synthesis* [14]; while all UMIs are supposed to be the same length (e.g. *k* = 12 bp), some miss several rounds of synthesis and end up as length 8. Then, due to the structure of 10x’s beads, the sequencer will keep reading into the 30-bp poly(dT) tail [18], mistakenly reading 4 Ts as the last 4 bases of the UMI. All genes follow the same trend, except for MALAT1, which we are able to identify and filter out (Section 4.2). Examining our 1k PBMC dataset, we compute the following position weight matrix characterizing the marginal probability distribution at each base pair of the UMI (Figure 2a).

**Fig. 2:**
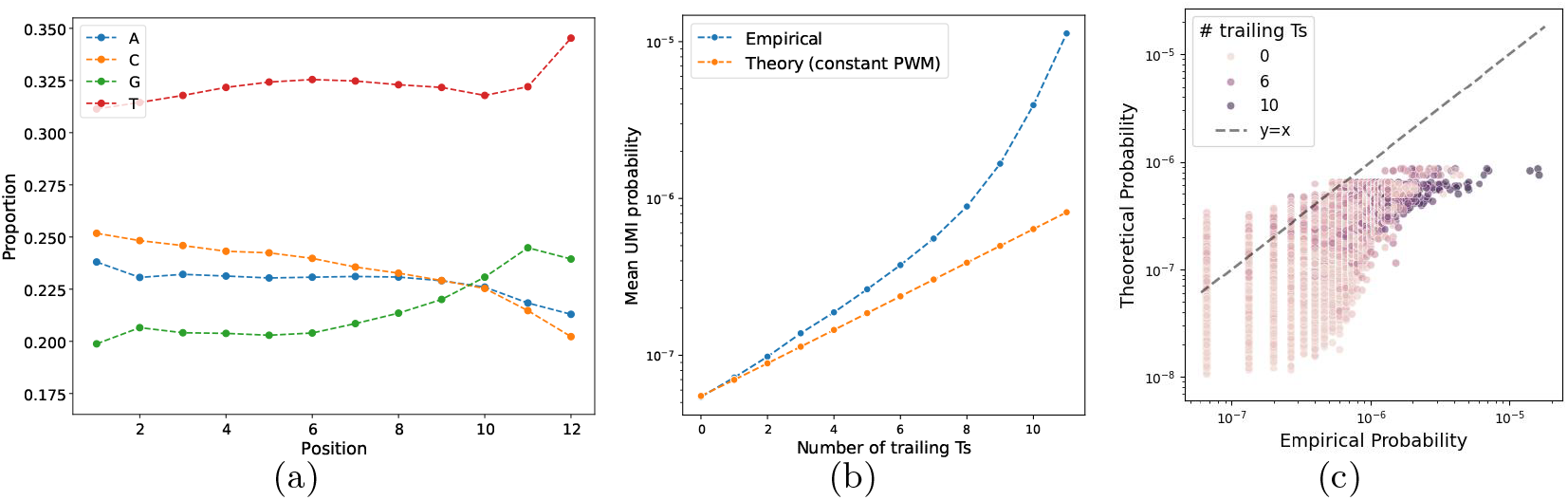
Analysis of empirical UMI nucleotide frequencies in 1k PBMC dataset. **a)** Position weight matrix (PWM) for the empirical UMI distribution across all genes except MALAT1. **b)** Mean probability for all UMIs given a certain number of trailing Ts. This is shown for the empirical frequencies of non-MALAT1 genes (blue), as well as the theoretical probabilities based on the constant PWM (Equation (8), orange). MALAT1 and PWM from **a)** are shown in Figure S6. **c)** Empirical UMI probabilities versus predicted probabilities using the constant PWM model in Equation (8).

The UMI frequencies can be well modeled by an independent, per-base probability distribution, which improves for shorter UMI lengths (Figure S1). For generating synthetic data with shortened UMI lengths, we truncate and only look at the first *k* base pairs of the UMI, as this mimics the true UMI synthesis process (discussed in Section S1.2). We model this probability by defining *n*_*L*_ as the number of occurrences of nucleotide *L* ∈ {*A, C, G, T*} in the UMI sequence *S*. Probabilistically, this independent model states that:

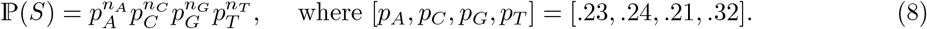

The per nucleotide probabilities are fit from the 1k PBMC dataset (Figure 2a), and hold stable across other 10x datasets with the same chemistry [14]. For non-10x chemistries, the specific probabilities may differ, but the same modeling approach (computing the empirical nucleotide frequencies and using Equation (8)) can be applied.

### 4.1 Failure in UMI synthesis

The only UMIs that significantly deviate from the model in Equation (8) are those that end with a string of Ts, as shown in Figure 2b,c. This is due to truncation during UMI synthesis, and with the 10x chemistry of 3^*′*^ sequencing, this leads us to read into the poly(dT) tail [14]. This is because of the specifics of 10x’s beads: to capture the mRNA transcript, each bead contains (in order) a TruSeq Read 1, the cell barcode (16-bp), the UMI (supposed to be 12-bp), poly(dT) sequence (30-bp), and finally VN (1-bp) an anchor to indicate the end of the poly(dT) stretch. However, in the sequential synthesis of UMIs, not all will make it to the full length 12. In fact, the authors claim that under 50% of 10x Chromium beads have the claimed length of 28 (16-bp barcode plus 12-bp UMI).

Mechanistically, at each iteration of UMI synthesis, there is some probability of failure to add the next base pair. In this case the UMI is capped at the end of that iteration, and no further base pairs are added. Assuming that the failure probability *q* is constant across all base pairs, this leads to a geometric distribution on the length of the synthesized UMI. When the UMI is shorter than 12-bp, the sequencer will read into the poly(dT) tail, leading to extra Ts at the end of the observed UMI. We show this process schematically in Figure S3, and refer the reader to [14] for further biochemistry details. Thus, we obtain the explicit probability of observing a UMI *S* with `T` trailing Ts by summing over all possible synthesis lengths *ℓ* from 12 − `T` to *k*.

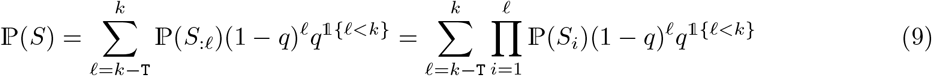

This model can be extended to allow for position-dependent failure probabilities *q*_*i*_, which when fitted to the data yield an extremely good fit (Figure 3c), details in Section S1.2. As these UMIs with many trailing Ts constitute a small fraction of all UMIs, we retain the simpler model in Equation (8) for computational and conceptual simplicity.

**Fig. 3:**
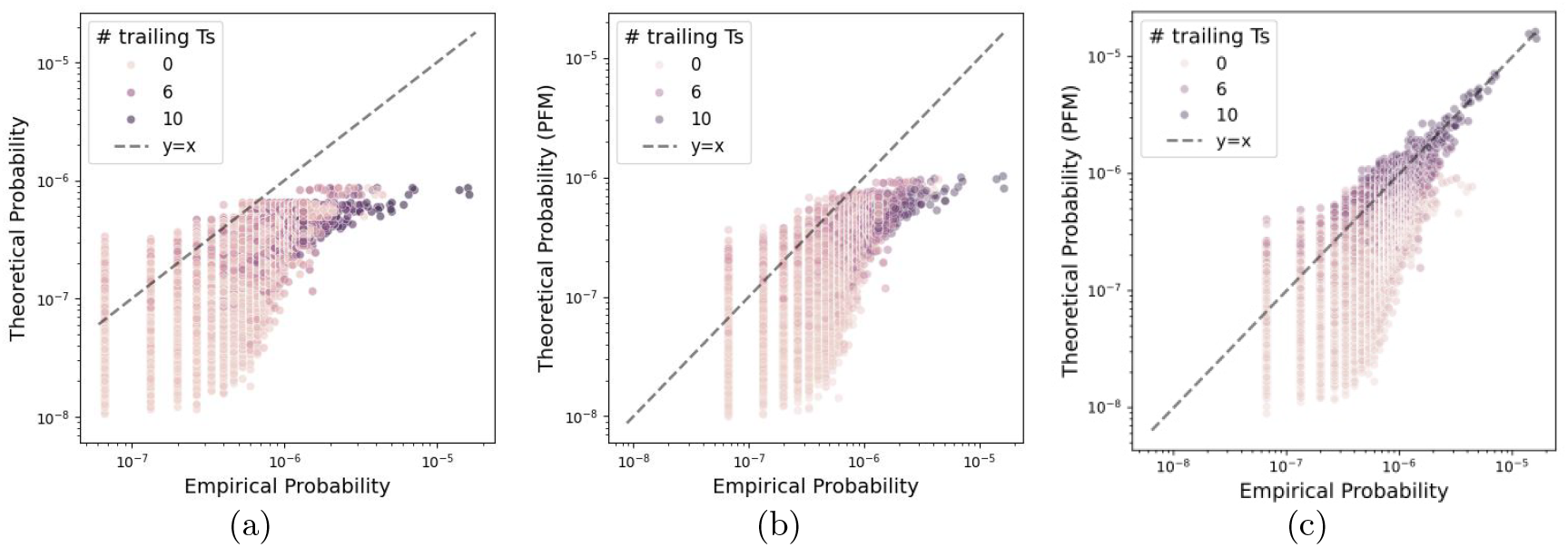
Fit of empirical 12-bp UMI probabilities using different models. **a)** Prediction using a constant PWM as in Equation (8). **b)** Prediction using PWM in Figure 2a. **c)** Prediction using truncated UMI synthesis model in Equation (9), monotonic per-bp synthesis probabilities fit as detailed in Section S1.2.

### 4.2 Poor fit of MALAT1

Analyzing the observed counts for shortened UMIs, our nonuniform model displays good accuracy across genes. However, we identify in this work that one gene, MALAT1, is a consistent outlier. Formally, studying the 1k PBMC dataset (Figure 4a), we observe that almost all counts fall within 3*σ* confidence intervals of their expectation under our model, except for MALAT1 (colored in red). As can be seen, these points follow a fundamentally different trend, and are negatively biased (i.e. experience more collisions). This same behavior was identified across all datasets studied.

**Fig. 4:**
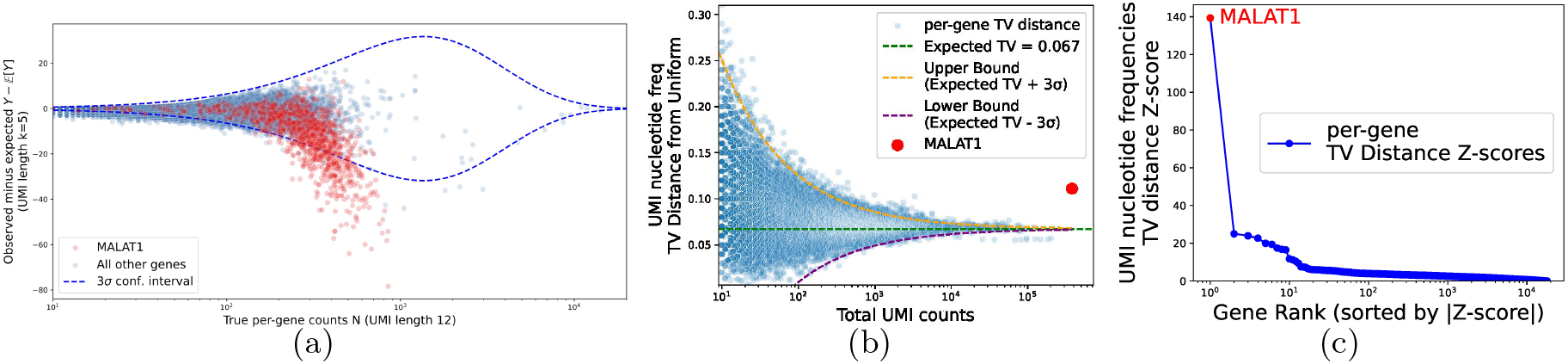
Outlier detection of MALAT1. **a)** 10x 1k PBMC dataset, corrected and deduplicated UMI counts. Nonuniform model (Equation (3)) with UMI frequencies generated by the marginal per-base pair model (Equation (8)). 3*σ* confidence intervals constructed from Equation (4) using nonuniform model. All genes aside from MALAT1 fall well within the 3*σ* confidence intervals. **b)** Per-gene analysis of TV distance between UMI nucleotide frequencies and mean nucleotide frequencies. Observed data is slightly overdispersed relative to the multinomial model, but MALAT1 is a clear outlier. **c)** Z-score computed for each gene’s TV distance, with variance approximation from Equation (12) (details in Section S1.5). Sorted z-scores validate the observation from **b)** that MALAT1 is a statistical outlier.

MALAT1, a long non-coding RNA, is notorious in the literature for its high expression, high rate of internal priming, and lack of polyadenylation [19, 20]. MALAT1 expression is commonly used for quality control [19], as it is retained within the cell nucleus, and so if high MALAT1 expression is not detected within a droplet, then the droplet is likely either empty or contains a poor-quality cell. MALAT1 does not have a poly(A) tail, but rather folds onto itself to form a unique and highly stable triple-helical A-rich structure [20].

We show that this deviation of MALAT1 can be detected de novo, directly from its UMI distribution (Figure 4). Concretely, in these plots we compute for each gene its empirical UMI distribution across all cells, and collapse this to a per-nucleotide distribution summed across all 12 positions. This yields 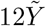 counts for 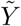 12-bp UMIs observed across all cells for this gene. After normalization, we compute the total variation (TV) distance between each gene’s per-nucleotide UMI distribution and the expected distribution (summing counts across all cells and all genes). Plotting this for all genes reveals MALAT1 as a clear outlier (Figure 4a). To provide confidence bands on the TV distance in Figure 4b, we formulate a simple null model that UMIs are i.i.d. from a common per-nucleotide distribution (details in Section S1.5). While reality is overdispersed relative to this simple null, it provides a useful framework for identifying outliers. Computing the z-score of each gene’s TV distance immediately identifies MALAT1 as the only gene that deviates significantly from the expected distribution (Figure 4c). Because of this, we exclude MALAT1 from further analyses: if desired, our method can be applied to MALAT1 using a separate MALAT1-specific UMI distribution. Because our model explicitly accounts for sequence composition, it can automatically flag biological anomalies like internal priming (MALAT1), acting as a quality control metric that naive counting may miss.

## 5 Estimator Validation

To validate the performance improvement afforded by our method, we empirically evaluated our collision-aware estimator on three publicly available 10x Genomics Peripheral Blood Mononuclear Cells (PBMC) datasets, of 1k (Figure 5) and 10k (Figure S7) cells sequenced using Chromium Single Cell 3^*′*^ v3 chemistry, and of 5k (Figure S8) cells sequenced using v4 chemistry. For each dataset, we artificially shortened the error-corrected length 12 UMIs to length *k* = 1, …, 12. For each UMI length, we utilized our estimator to predict the true collision-free counts from the number of unique length *k* UMIs observed (details in Section S1). As the ground truth, we used the counts obtained from the full length 12 UMIs, which are largely collision-free; for a uniform UMI distribution, the expected number of unique length 12 UMIs (from Equation (5)) observed for *N* = 1000 is *Y* = 999.97, and for the maximum observed count of highly expressed genes with *N* = 10000 is *Y* = 9997. To evaluate the performance of our estimator and its collision-oblivious counterpart (which estimates 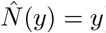), we measured their raw gene expression estimation accuracy, as well as their performance in downstream differential expression (DE) analysis.

**Fig. 5:**
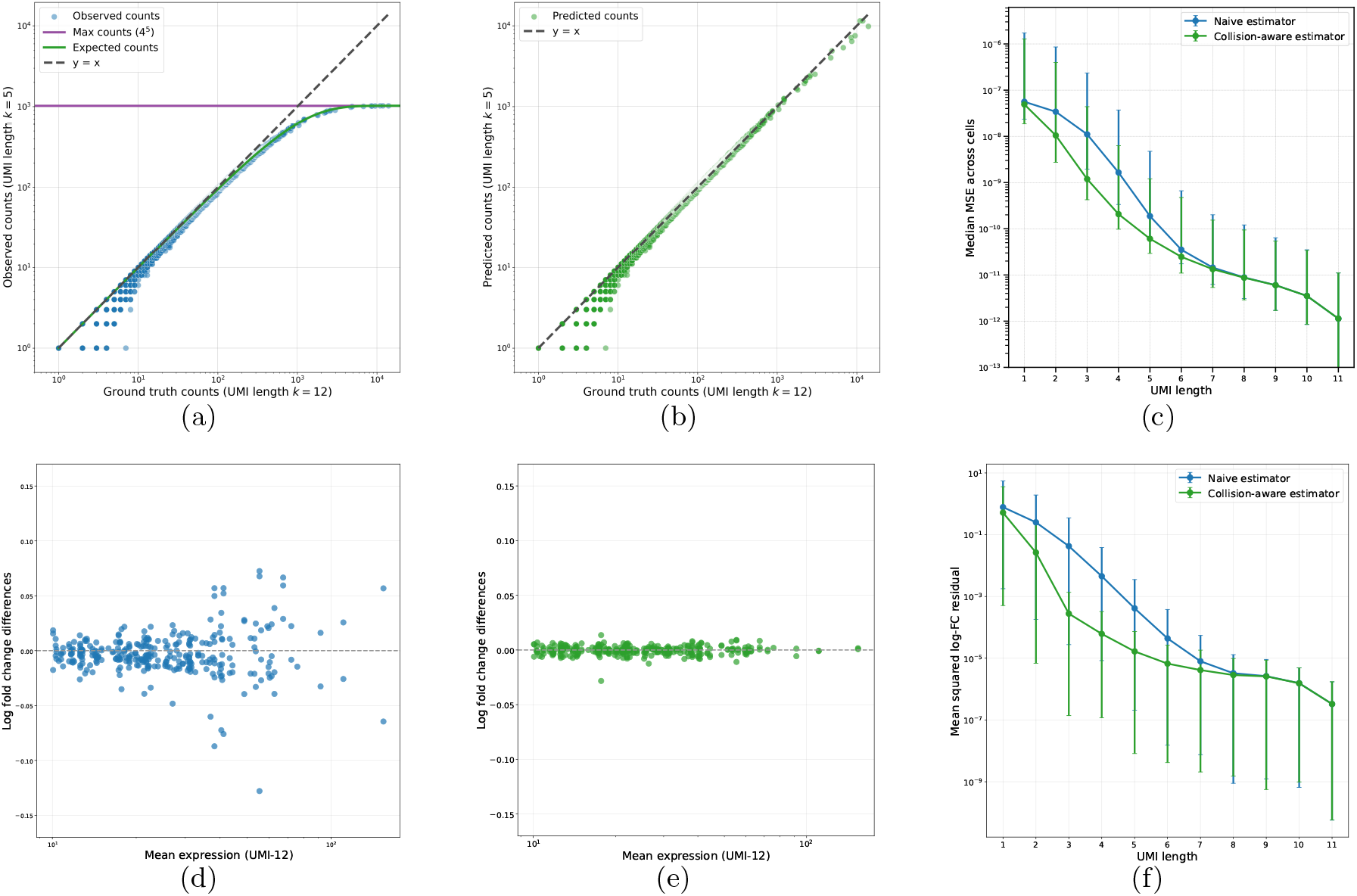
Improved performance of method-of-moments estimator on 10x’s PBMC 1k dataset. Panels **a-c** demonstrate enhanced accuracy in raw expression estimation, while panels **d-f** show the downstream improvement for log-fold change (LFC) estimation of differentially expressed genes (all genes shown in Figure S11). Processing details in Section S1. **a)** Concordance between our statistical model of collisions and the observed data for a UMI length of *k* = 5. **b)** Same as **a** but using our collision-aware estimator. MSE is reduced by 95% from **a** to **b. c)** Per-cell MSE between normalized gene expression vectors estimated from shortened UMIs and the ground truth (k=12), shown as a function of UMI length (median and 95% CIs). **d)** Difference between predicted (using *k* = 5, naive estimator) and true LFC (predicted - ground truth). **e)** Same as **d** but using our collision-aware estimator. MSE is reduced by 96% from **d** to **e. f)** MSE for estimating LFC (as in **d**,**e**), evaluated across UMI lengths (95% CIs).

We see the dramatic impact of collisions at shorter UMI lengths like *k* = 5 (Figure 5a), which our method-of-moments estimator is able to correct for (Figure 5b). To aggregate this metric across genes within a cell, we normalize the gene expression vectors and compute the MSE between our estimates and the ground truth *k* = 12 gene expression. Computed across all cells, our estimator reduced the mean squared error (MSE) by 95% over the naive estimator at a UMI length of 5 (Figure 5c). These performance improvements generalize to both additional datasets (Section S2).

Our more accurate quantification translates to improved retention of biological insights in downstream tasks. For the prototypical task of cell type prediction, we observe that even with very short UMIs (e.g. *k* = 4), the naive estimator still allows accurate CellTypist classification [21] (over 90% accuracy, Figure S9). To generate a more meaningful comparison, we focus on differential gene expression analysis. For low-expression marker genes, presence/absence of this gene is often sufficient, and so UMI length has little effect (Figure S10). Studying more highly expressed genes, we show that our improved quantification ensures accurate estimation of their fold changes (Figure 5d,e). Aggregating the error in fold change estimation across genes highlights the improved performance of our estimator below *k* = 8; at *k* = 5, our estimator reduces MSE by 96% (Figure 5f).

Theoretically, our estimator yields dramatic performance improvements. We highlight these in Table 1, comparing the bias, variance, and resulting MSE of our collision-aware estimator to the naive estimator across different regimes of the number of unique transcripts *N* relative to the number of unique UMIs *K*. We assume a uniform UMI distribution for simplicity. Detailed derivations are provided in Section S7.1. Throughout, we use standard Bachmann–Landau (big-O) asymptotic notation. Our results show that in the no-collision regime 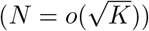, both estimators perform well.

However, in the moderate-collision regime (*N* = *o*(*K*), 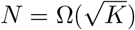), our estimator remains unbiased with no excess variance beyond that present in *Y*, while the naive estimator is bias-dominated with MSE scaling as Θ(*N* ^4^*/K*^2^). In the high-collision regime (*N* = *o*(*K* log *K*), *N* = Ω(*K*)), our estimator begins to show increasing bias and variance inflation, but still outperforms the linear in *N* bias of the naive estimator. Finally, in the saturation regime (*N* ≥ *cK* log *K* for *c >* 1), both estimators are fundamentally limited by the lack of information in *Y*, and exhibit linear in *N* bias. We further show in Sections S3 and S4 that the Poissonized maximum likelihood estimator, while theoretically optimal in the nonuniform setting, provides negligible practical improvement over the method-of-moments estimator, validating the latter as the recommended approach.

## 6 Discussion

In this work we showed that the ubiquitous approach of collapsing reads with identical UMIs as PCR duplicates is statistically biased. While conventionally the solution to this is to use longer UMIs to minimize the number of collisions, we enable the use of shorter and cheaper UMIs via a method-of-moments estimator that adjusts for these UMI collisions. We show that our estimator is near-optimal theoretically, matching the empirical performance of the poissonized MLE, and that in practice it performs extremely well up until the threshold of saturation. This allows for robust biological and statistical conclusions to be drawn from UMIs as short as 5 bp, closely matching the results from the standard 12-bp UMIs. This has the potential to further reduce sequencing costs and simplify manual UMI synthesis, enabling more cost-effective and scalable sequencing. There are many exciting directions of future work, including incorporating error correction into our statistical model, and leveraging per-UMI counts as opposed to just the presence/absence of a given UMI.

## 7 Declarations

### 7.1 Ethics approval and consent to participate

Not applicable.

### 7.2 Consent for publication

Not applicable.

### 7.3 Availability of data and materials

All datasets analyzed in this study are publicly available from 10x Genomics (PBMC 1k, 5k, and 10k datasets):

- 1k PBMC: https://www.10xgenomics.com/datasets/1-k-pbm-cs-from-a-healthy-donor-v-3-chemistry-3-standard-3-0
- 10k PBMC: https://www.10xgenomics.com/datasets/10k-human-pbmcs-3-v3-1-chromium-controller-3-1-high
- 5k PBMC: https://www.10xgenomics.com/datasets/5k_Human_Donor1_PBMC_3p_gem-x

Source code implementing the collision-aware estimator is available at: https://github.com/agdylan/minimal_UMIs under the MIT License. The version used for this manuscript corresponds to commit id: 021a029.

### 7.4 Competing interests

The authors declare that they have no competing interests.

### 7.5 Funding

D.A. was supported by R25HG006682. RAI was supported in part by funding from NIH Grants R35GM131802, and R01HG005220. T.Z.B. was supported by the Eric and Wendy Schmidt Center at the Broad Institute.

### 7.6 Authors’ contributions

DA implemented the collision-aware estimator, carried out analyses, and drafted the manuscript. RAI provided guidance throughout the project and contributed to manuscript revisions. TZB conceived and supervised the study, performed analyses, and drafted the manuscript. All authors read and approved the final manuscript.

## 7.7 Acknowledgments

We thank Elvira Forte for her careful review of the manuscript and for insightful suggestions that improved its clarity.

## 7.8 Use of large language models

Large language models (LLMs) were used to assist in modifying / writing boilerplate code, and in generating plotting scripts. All computational analyses, algorithms, and results reported in the manuscript were authored, executed, and validated by the authors. Similarly, LLMs were used for minor language editing (grammar/clarity) of author-written text. All text was reviewed and approved by the authors.

## Supplementary Information

### Supplement roadmap

This supplement provides additional methodological detail, extended theoretical results, and supplementary analyses supporting the main text. Section S1 details data processing, synthetic UMI truncation, fitting the truncated-synthesis model, sequencing-error considerations, differential expression analysis details, and MALAT1 analysis details. Section S2 provides analogues of our 1k PBMC results figure (Fig. 5) on two additional PBMC datasets (10k v3 and 5k v4) to validate the generality of our method, and includes DE diagnostics. Section S3 derives the maximum likelihood estimator under a Poissonized model and analyzes its properties. Section S4 empirically validates different UMI distribution models and collision-aware estimators, showing that the method-of-moments estimator with a constant PWM captures nearly all available gains. Section S5 derives estimator variance and convexity (and the *Y* = *K* extrapolation). Section S6 leverages the asymptotic normality of *Y* to characterize the bias and variance of our estimator. Section S7 computes and compares the MSE of our estimator and the naive estimator across different regimes of *N* (details for Table 1). It also proves the impossibility of estimation for *N* ≥ *cK* log *K* for *c >* 1, and shows that our estimator matches the Cramér–Rao lower bound in a simplified binomial setting.

## S1 Datasets and processing details

We processed all datasets using Cell Ranger. According to Cell Ranger, the 1K PBMC (v3) dataset contains 1222 cells sequenced at an average depth of 54k reads/cell, the 10K PBMC dataset (v3) contains 11485 cells sequenced at an average depth of 30k reads/cell, and the 5k PBMC dataset (v4) contains 4782 cells sequenced at an average depth of 39k reads/cell. To synthetically generate shortened UMIs, we processed the output BAM file from Cell Ranger, and shortened the UB field (corrected UMI, discussed in Section S1.3) to the first *k* base pairs for UMI length *k* (removing the last 12-*k* base pairs)

For downstream analyses, for each dataset and UMI length we performed the standard pipeline of library-size (total count) normalization, scaling to 10,000 counts per cell followed by a log1p transformation. We use CellTypist [21] to annotate each cell, using their `Healthy_COVID19_PBMC` model, which is trained on “peripheral blood mononuclear cell types from healthy and COVID-19 individuals”, as this is the best match for our datasets.

### S1.1 Synthetically truncated UMIs

**Fig. S1:**
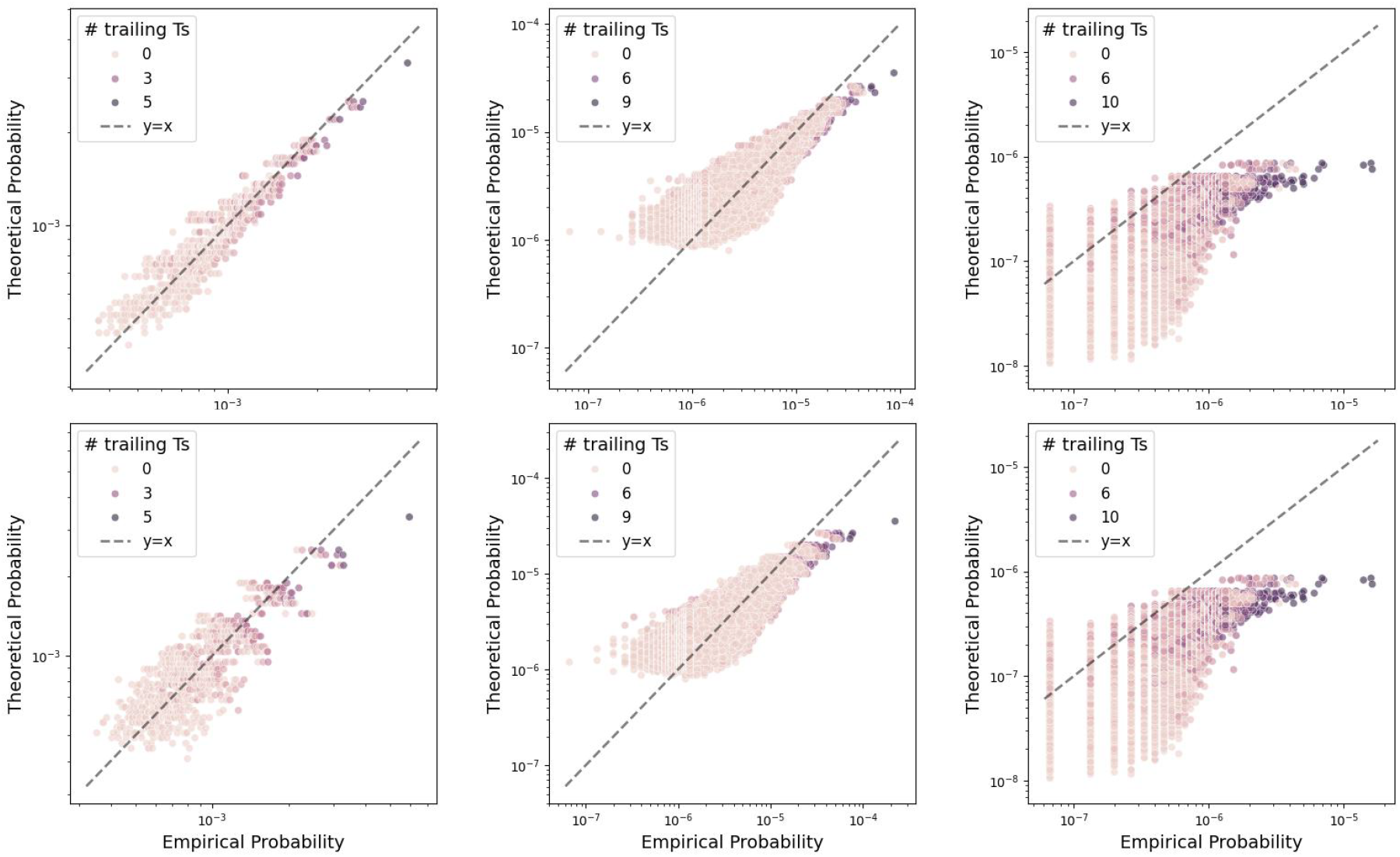
Accuracy of UMI frequency model in Equation (8). Columns correspond to UMI lengths of 5 (left), 9 (center), and 12 (right). Top row corresponds to truncating UMIs from the rear (removing the last 12 − *k* base pairs, as we use for all datasets), and the bottom row to truncating UMIs from the front (to show our model’s robustness). Note that the last column is the same for both rows, as the UMIs are full length (this plot is also included in Figure 3a). Both settings are fit by the constant PWM model reasonably well, with an improved fit when *k* is short.

### S1.2 Fitting truncated UMI synthesis model

UMI synthesis is an error-prone procedure. Recall the truncated UMI synthesis model posited in Equation (9) for a UMI *S* of length *k* with `T` trailing Ts, motivated by [14]:

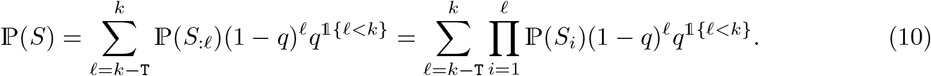

This is based on the assumption that each base pair is synthesized successfully with probability 1 − *q*, independently across base pairs. When synthesis fails at a given base pair, the UMI is capped at that position (length *< k*), and the sequencer reads past the end of the truncated UMI into the poly(dT) tail, leading to trailing Ts in the observed UMI (shown schematically in Figure S3).

We see that this model can be extended to the case of per-bp synthesis failure probabilities *q*_*i*_:

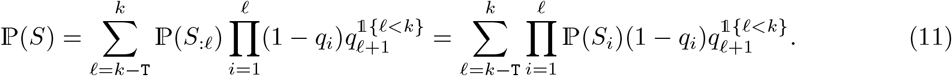

We fit this model for the aggregated statistics over values of `T` in the 1k PBMC dataset, using the nucleotide frequency from the first 6-bp: [.23, .25, .20, .32] (as opposed to [.23, .24, .21, .32] when computed over the full UMI). We show that fitting monotonically increasing *q*_*i*_ provides a good balance between flexibility and avoiding overfitting, as shown in Figure S2. The monotonically increasing *q*_*i*_ fitted on the 1k PBMC dataset yield good predictive performance for the 10k PBMC dataset, as shown in Figure S4.

**Fig. S2:**
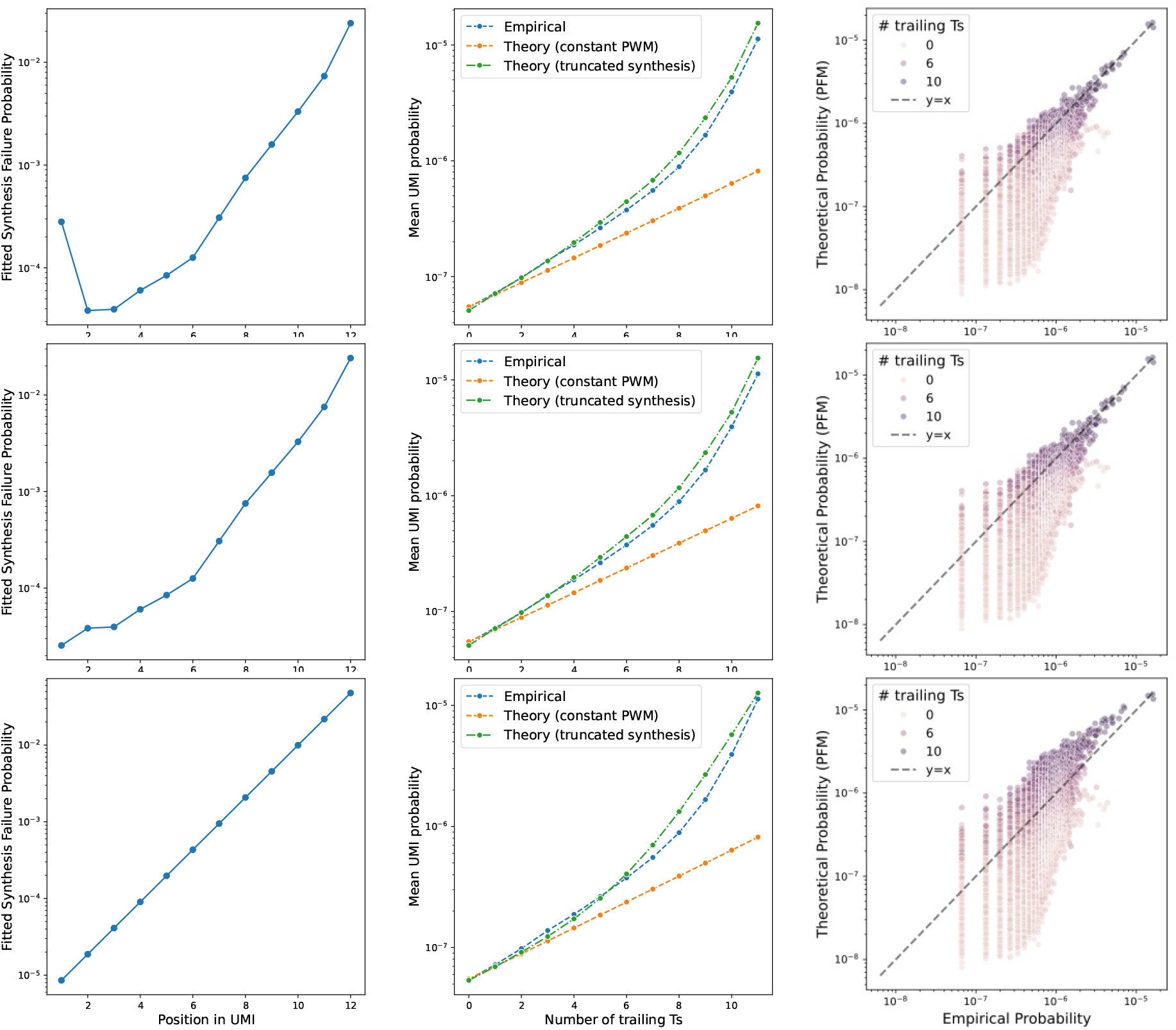
Fitting per-bp UMI synthesis failure probabilities. We fit three models for the per-bp synthesis failure probabilities *q*_*i*_, where the first row corresponds to unconstrained *q*_*i*_, the second row to monotonically increasing *q*_*i*_, and the third row to linearly increasing *q*_*i*_ in log-scale. The left column shows the fitted *q*_*i*_, the center column the mean observed vs. predicted UMI probabilities grouped by the number of trailing Ts, and the right column the observed vs. predicted UMI probabilities on a per-UMI basis.

**Fig. S3:**
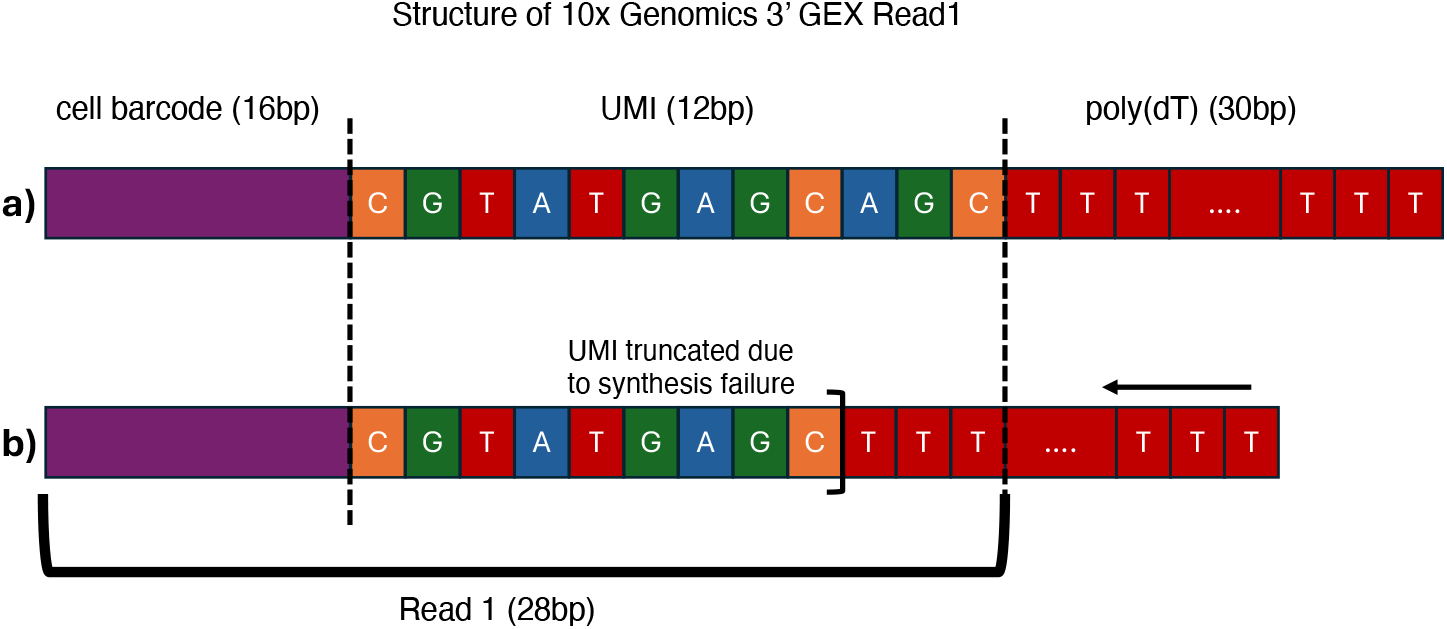
Schematic of Read1, how synthesis failure leads to trailing Ts. During library preparation, a poly(dT) tail is synthesized immediately after the UMI. If UMI synthesis fails at a given base pair, the sequencer reads past the end of the truncated UMI into the poly(dT) tail, leading to trailing Ts in the observed UMI. **a)** shows successful synthesis of a full length UMI, where the 16bp barcode is fully present, and the 12bp UMI is fully synthesized, leading to the desired 28bp Read1. **b)** shows a UMI where synthesis fails at the 10th base pair, leading to a truncated UMI of length 9. However, the sequencer reads past the end of this truncated UMI, leading to 3 trailing Ts.

**Fig. S4:**
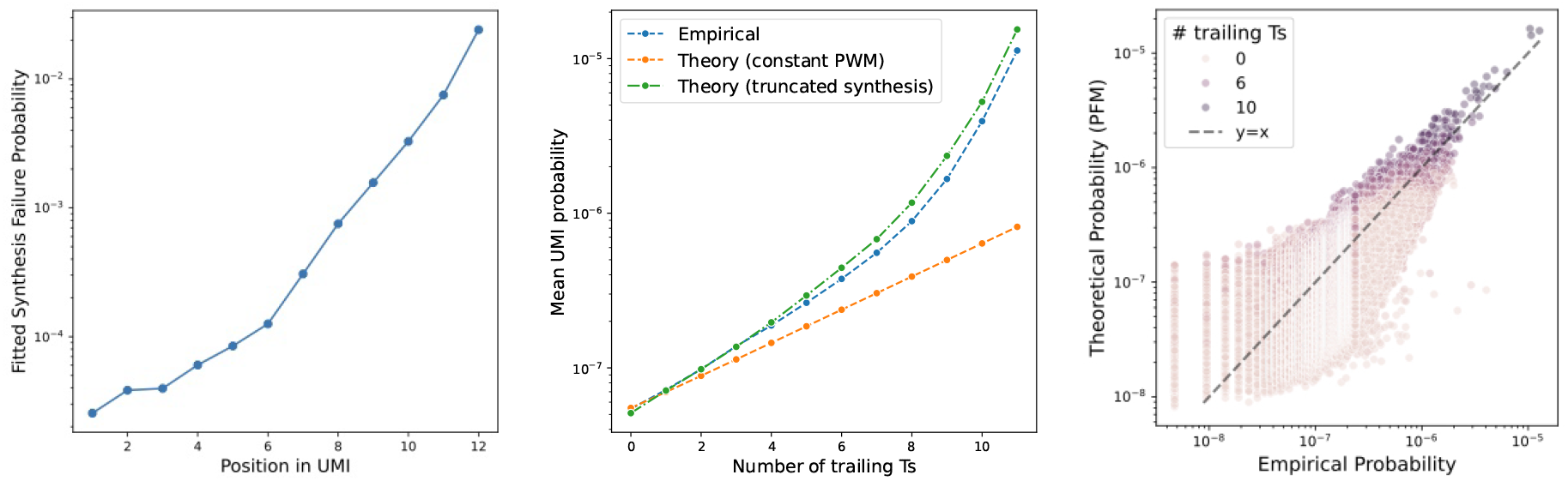
Evaluating per-bp UMI synthesis failure probabilities on 10k PBMC dataset. We leverage the fitted monotonically increasing *q*_*i*_ from the 1k PBMC dataset (left) to predict UMI frequencies in the 10k PBMC dataset. The center column shows the mean observed vs. predicted UMI probabilities grouped by the number of trailing Ts, and the right column the observed vs. predicted UMI probabilities on a per-UMI basis. Due to the larger dataset size, and hence more low count UMIs observed, the fit is not as close as in the 1k PBMC dataset, but the overall trends are still captured well. A more careful fitting procedure, better taking into account the absent UMIs could yield an improved / generalizable fit.

### S1.3 Sequencing error

As noted, our model does not account for UMI sequencing errors. Errors introduced during PCR amplification or during sequencing impact the actual UMIs we observe, leading to incorrectly undeduplicated counts: UMIs that should have been deduplicated but are not due to errors. Consider for concreteness the case where a gene has 100 reads associated with it, all stemming from 2 unique UMIs. However, during sequencing, one of these reads suffers a sequencing error at the last base pair of its UMI. Then, naively, the number of unique UMIs recorded as the ground truth for this gene is 3. However, for a UMI length of 11 or shorter, if we only observe the first *k* base pairs of the UMI, our count will now be only 2, as the errored base pair will be deleted from the UMI and there will only be the true 2 unique UMIs.

For simplicity, in this work we analyze Cell Ranger’s error corrected UMIs (UB instead of UR field in BAM file), noting that these are functions of the full length UMI which we would in practice not observe. Algorithmically, UMI error correction methods utilize the fact that neighboring UMIs in Hamming space (i.e. UMIs one base pair apart) are very rare naturally for long UMIs, and so these are collapsed and considered sequencing errors. However, as the UMI length shortens, the number of possible UMIs decreases exponentially, and so the probability of two UMIs being one base pair apart increases. In fact, a naive use of existing error correction methods, e.g. collapsing UMIs which are Hamming distance 1 and one has at least a factor of 2 plus 1 more counts than the other [11], may yield a *non-monotonic* relationship between the observed UMI counts and the true UMI counts, as the UMI space becomes so saturated that many valid UMIs are incorrectly collapsed together.

This issue of UMI error correction for short UMIs was briefly studied in [13], with a Bayesian method that worked to jointly model collisions and sequencing error, but due to the computationally intensive nature of their proposed estimator their approximate solution had to be further simplified by quantizing a dynamic programming problem. Note that any UMI error correction methods can be used as preprocessing for our algorithm, which can work to avoid such issues by utilizing information such as Phred scores, counts, and graph-based rules [10–12].

**Fig. S5:**
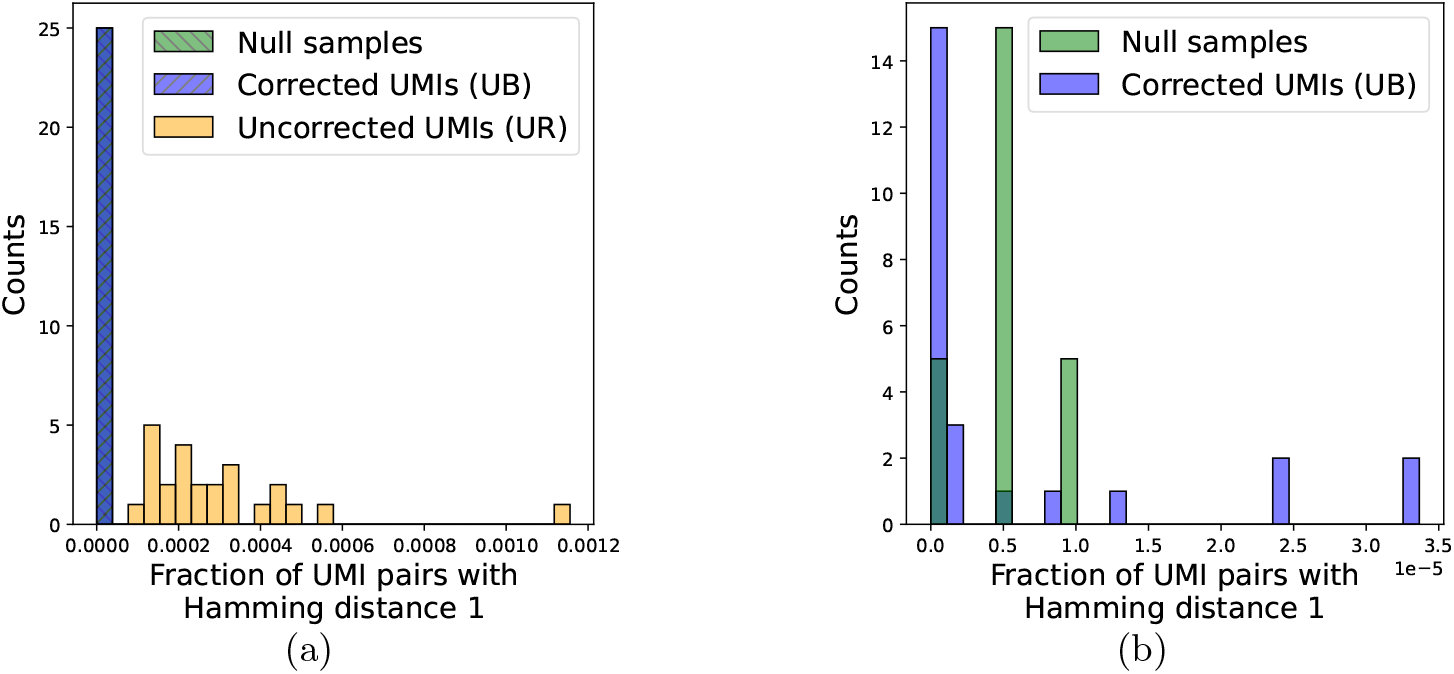
Pairwise Hamming distance between UMIs shows errors in uncorrected UMIs. We compute pairwise Hamming distances between all UMIs for a given gene in a given cell. Null samples generated by sampling *n* UMIs without replacement from the empirical UMI distribution across all cells all genes. We plot the top 5 genes over the top 5 cells, and choose *n* as the mean UB counts over these. **a)** shows all three methods, and **b)** zooms in on the corrected UMIs and null samples.

An empirical analysis of the 1k PBMC dataset reveals that many more uncorrected UMIs are 1 base pair apart than would be expected by chance (Figure S5). We study the top *m* cells, and their expression of the top *m* most expressed genes (*m* = 5). We compute, across all pairs of UMIs, what fraction are 1 base pair apart. In addition to the uncorrected (UR) and corrected (UB) UMIs, we also generate a null distribution by randomly sampling a matching number of UMIs from the empirical error corrected UMI distribution (UB) across all cells and all genes. Plotting histograms of these three, we see that the uncorrected UMIs have a significantly higher fraction of pairs that are 1 base pair apart compared to the corrected UMIs and the null distribution, separated by an order of magnitude (Figure S5a). Zooming in on the corrected UMIs and the null samples, these are statistically very similar, validating our theoretical model that UMIs within a given cell, for a given gene, are drawn i.i.d. from some common distribution across cells / genes (Figure S5b).

### S1.4 Differential Expression processing details

We perform differential expression analysis to assess our estimator’s ability to recover biological insights. To this end, we select cell types with over 100 cells in the 1k dataset (CD14 monocytes, Naive B cells, and Naive CD4+ T cells) and run the DE analysis pipeline as described below on each of them, and aggregate the results. For each counts matrix, after normalization and log1p transformation, we run scanpy’s rank genes group function with the Wilcoxon rank sum test to compute differential expression statistics [22]. For each gene we compute the difference between the estimated and ground truth log-fold changes, using either the naive estimator or collision-aware one. An example of this is shown in Figure 5d-f.

In our analysis, we observe several marker genes for which UMIs are unnecessary for detecting differential expression. For example, we observe that FCN1 is a marker gene for CD14 monocytes, which displays essentially a binary expression pattern (Figure S10, additional discussion in Section S2.1). Simply the presence or absence of this gene (UMI length of 0) is sufficient. Such genes are not of interest to study here, as UMIs are not even needed in the first place. To this end, we filter for genes with mean expression greater than 10 across all cells in the dataset. Additionally, similar to a volcano plot, we filter for genes with at least a 25% fold change in either direction, with an adjusted p-value less than 0.05.

**Fig. S6:**
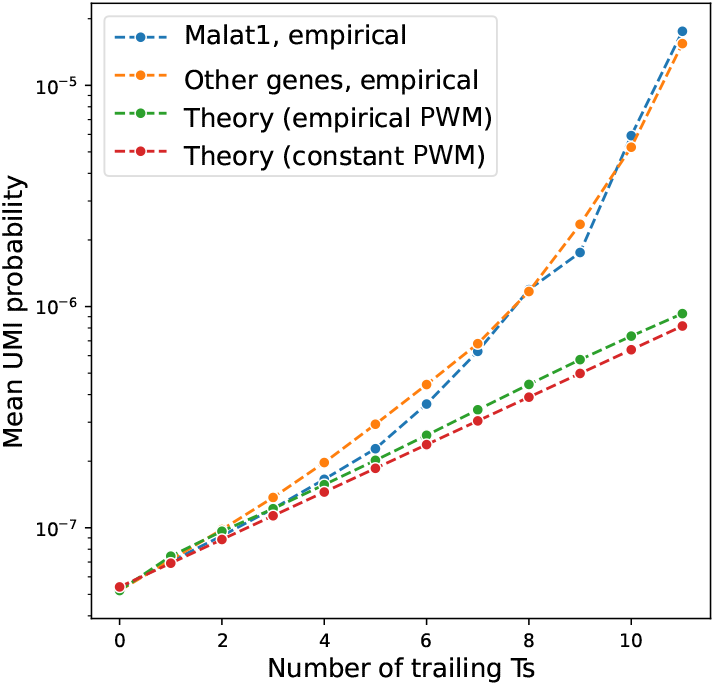
Analysis of empirical UMI nucleotide frequencies in 1k PBMC dataset. Mean probability for all UMIs given a certain number of trailing Ts. This is shown for the empirical frequencies of MALAT1 (blue) and all other genes (orange), as well as the theoretical probabilities based on the PWM (subfigure **a)**, green) and the constant PWM (Equation (8), red).

### S1.5 MALAT1 analysis

Here, we provide the details for the UMI-based identification that MALAT1 is an outlier (shown in Figure 4) We can approximate the variance of the TV distance for *m* counts by noting that under the model that the nucleotides of a UMI are independent and identically distributed, we are computing the variance of the TV distance between a sample of size *m* from a multinomial and its expectation. For large *m*, the entries of the multinomial are approximately independent. Concretely, denoting ***p*** as the nucleotide distribution, and *X* ∼ Multinomial(*m*, ***p***):

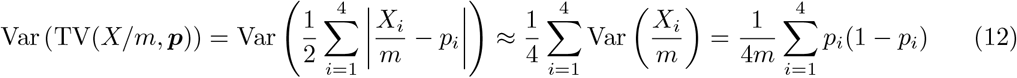

### S1.6 Synthetic data generation for Figure 1

For Figure 1, we generate synthetic data to show the natural behavior when UMIs are long (Figure 1b), when they are short and yield collisions (Figure 1d), and how we can correct for this by utilizing our method-of-moments estimator (Figure 1e). To generate the synthetic UMI data, we simulated the labeling process under a uniform distribution. For a given UMI length *k*, UMIs are simulated as being drawn uniformly at random with replacement from all *K* = 4^*k*^ possible sequences. To model a true transcript abundance of *N*, we take *N* independent draws from this pool and record the resulting number of unique UMIs observed as *Y* . This process was evaluated across logarithmically spaced values of *N* to capture the system’s behavior from a low-collision regime up through complete saturation.

## S2 Validation on additional datasets

Throughout, we discussed the application of our method to 10x’s PBMC 1k dataset. Here, we show that our method’s performance improvements hold in general. We recapitulate our analyses from Figure 5 for the 10x Genomics 10k PBMC dataset with v3 chemistry (Figure S7), and for the 10x Genomics 5k PBMC dataset with v4 chemistry (Figure S8).

**Fig. S7:**
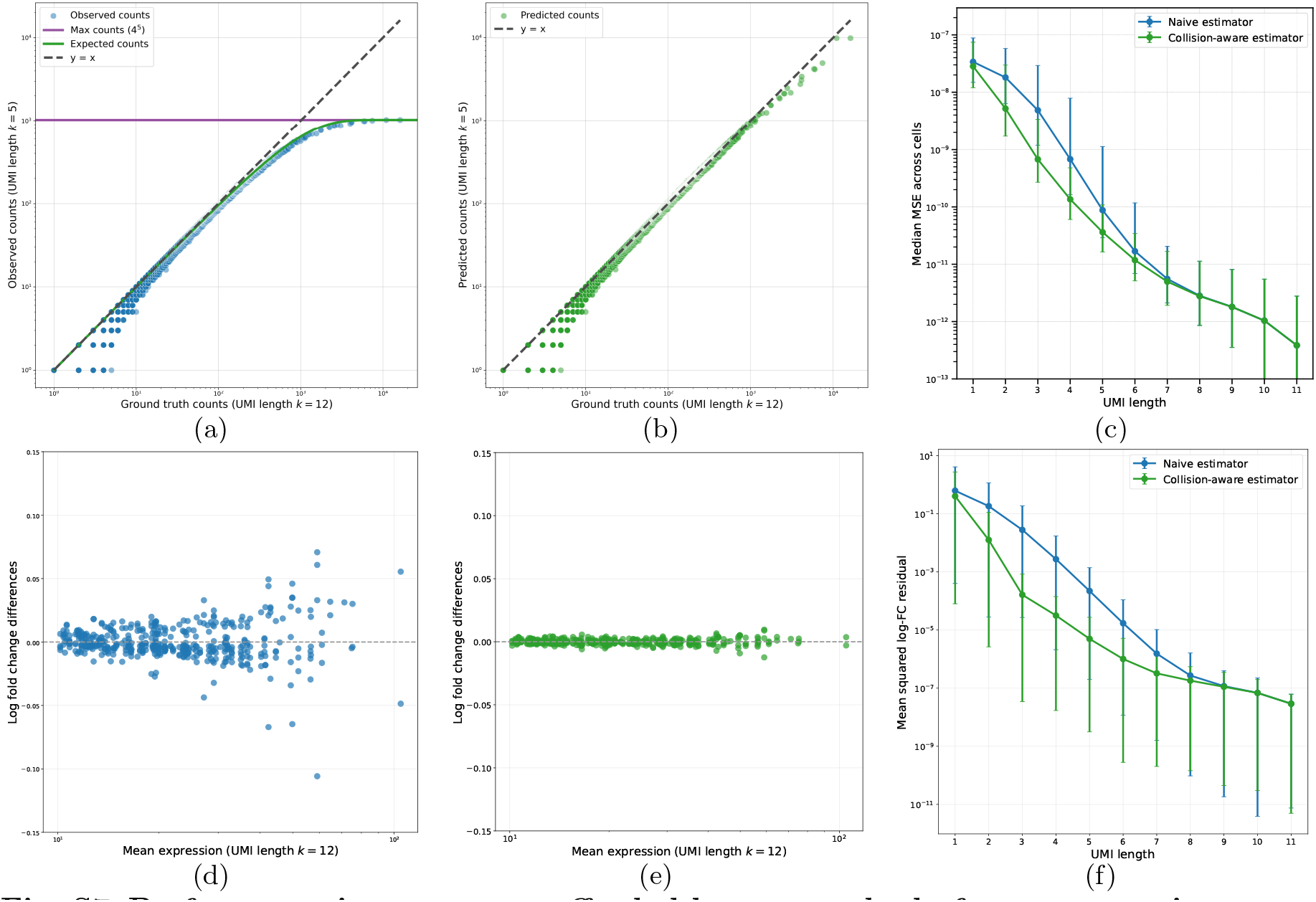
Performance improvement afforded by our method-of-moments estimator on 10x’s PBMC 10K dataset (v3 chemistry). **a-c** show improvement in raw expression estimation, and **d-f** show improvement for log-fold change (LFC) estimation of differentially expressed genes, replication of Figure 5.

**Fig. S8:**
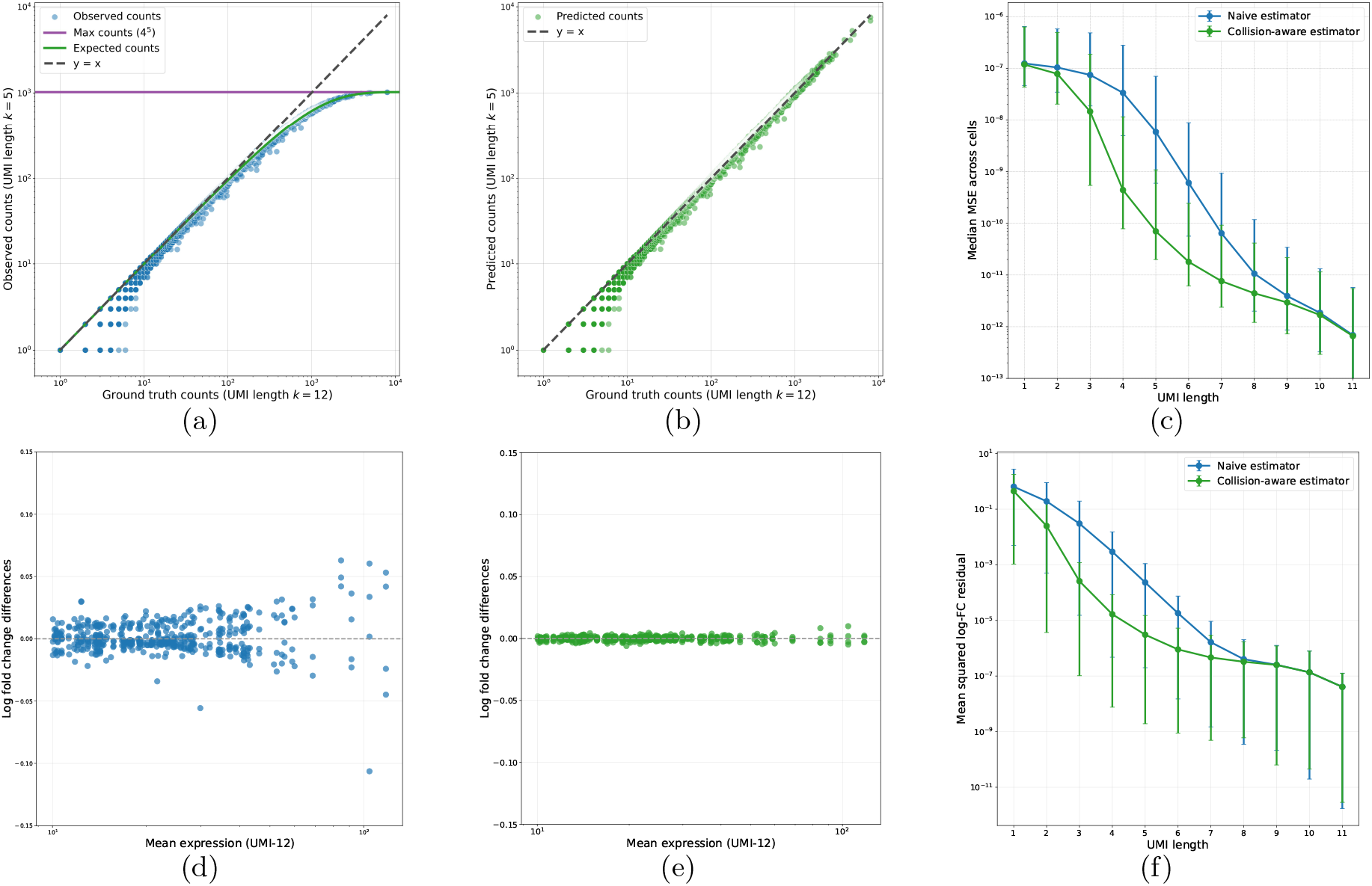
Performance improvement afforded by our method-of-moments estimator on 10x’s PBMC 5K dataset (v4 chemistry). **a-c** show improvement in raw expression estimation, and **d-f** show improvement for log-fold change (LFC) estimation of differentially expressed genes, replication of Figure 5.

### S2.1 DE supplemental figures

To begin, in Figure S9, we highlight the ease of cell type annotation as a computational task. Even for very short UMI lengths like *k* = 4, we still retain essentially the same accuracy with the naive estimator as for *k* = 9, showing the cell type annotation is insufficiently sensitive to actual counts (and by extension, UMI length) to serve as our benchmark.

**Fig. S9:**
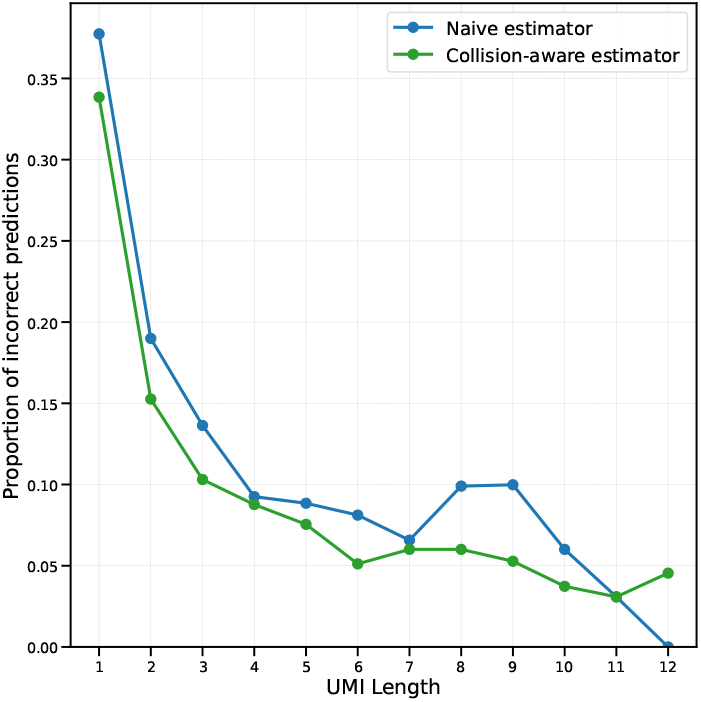
Fraction of errors in cell type prediction using CellTypist as a function of UMI length.

As discussed in Section S1.4, we filter out low count differently expressed genes from our analysis. This is because, for certain marker genes that exhibit a binary-like expression (0 in certain cells, and nonzero in others), we don’t actually need UMIs *at all* to detect that this gene is differentially expressed. FCN1, a marker gene for CD14 Monocytes, overwhelmingly displays 0 expression for other cell types, with only 12% of CD14 monocytes having 0 expression, while over 93% of other cells have 0 counts (Figure S10). With a UMI length of 12, the LFC is 5.9 with a p-value of 1.1 × 10^−118^, while with a UMI length of 0 the naive estimator provides an LFC of 4.2 with a p-value of 2.5 × 10^−110^. Clearly, an improved estimator is unnecessary in this situation, and so we filter out genes with low average expression to ensure that we are only comparing those where UMI length will play a role.

**Fig. S10:**
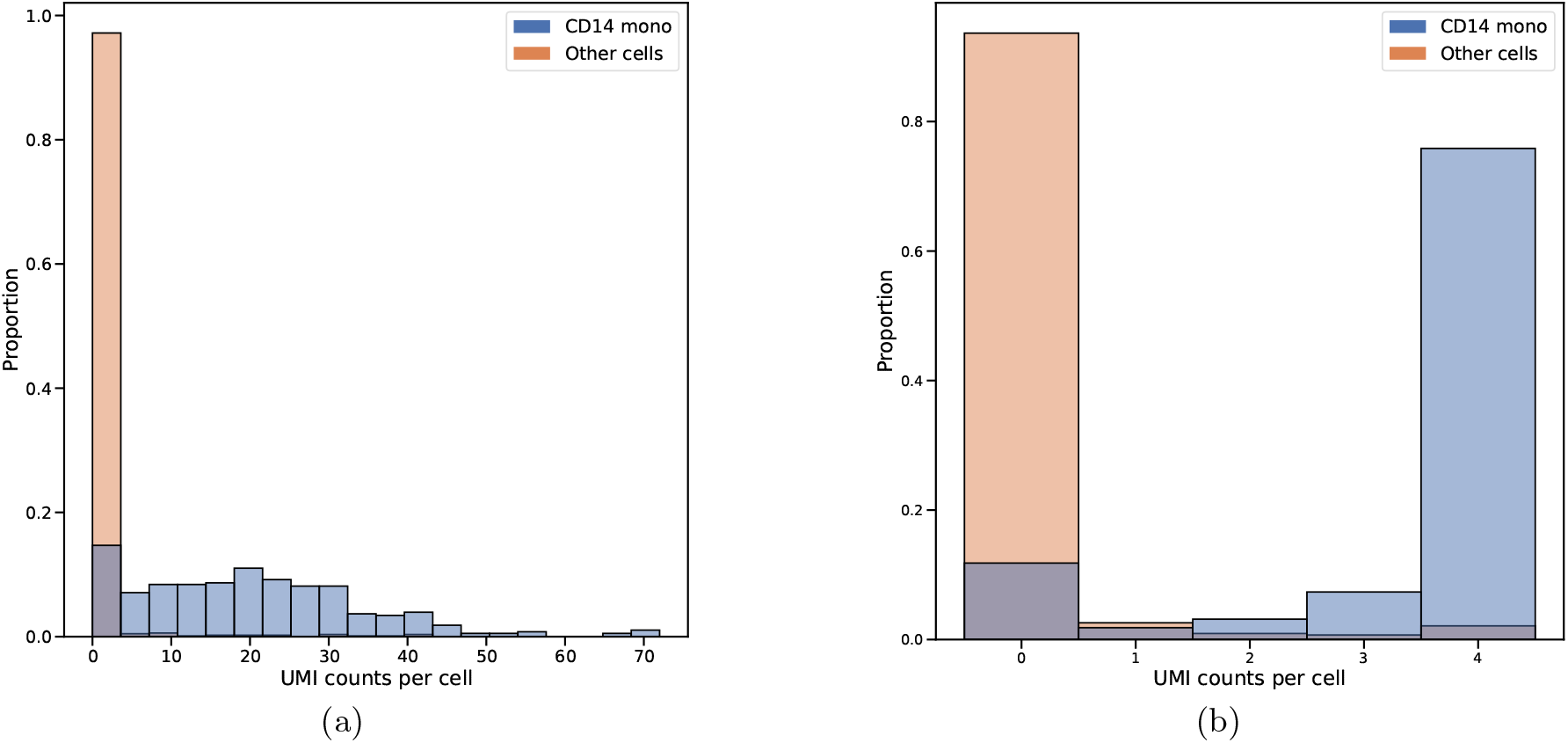
Marker genes do not require UMIs to be called as DE. Marker genes that display a binary expression pattern are called as differentially expressed even by the naive estimator for a UMI length of 1. Shown is FCN1, a marker gene for CD14 monocytes. **a)** Distribution of raw UMI counts for the gene FCN1 analyzing the full length 12 UMIs. **b)** Same as **a** but using the raw length 1 UMI counts (the naive estimator).

**Fig. S11:**
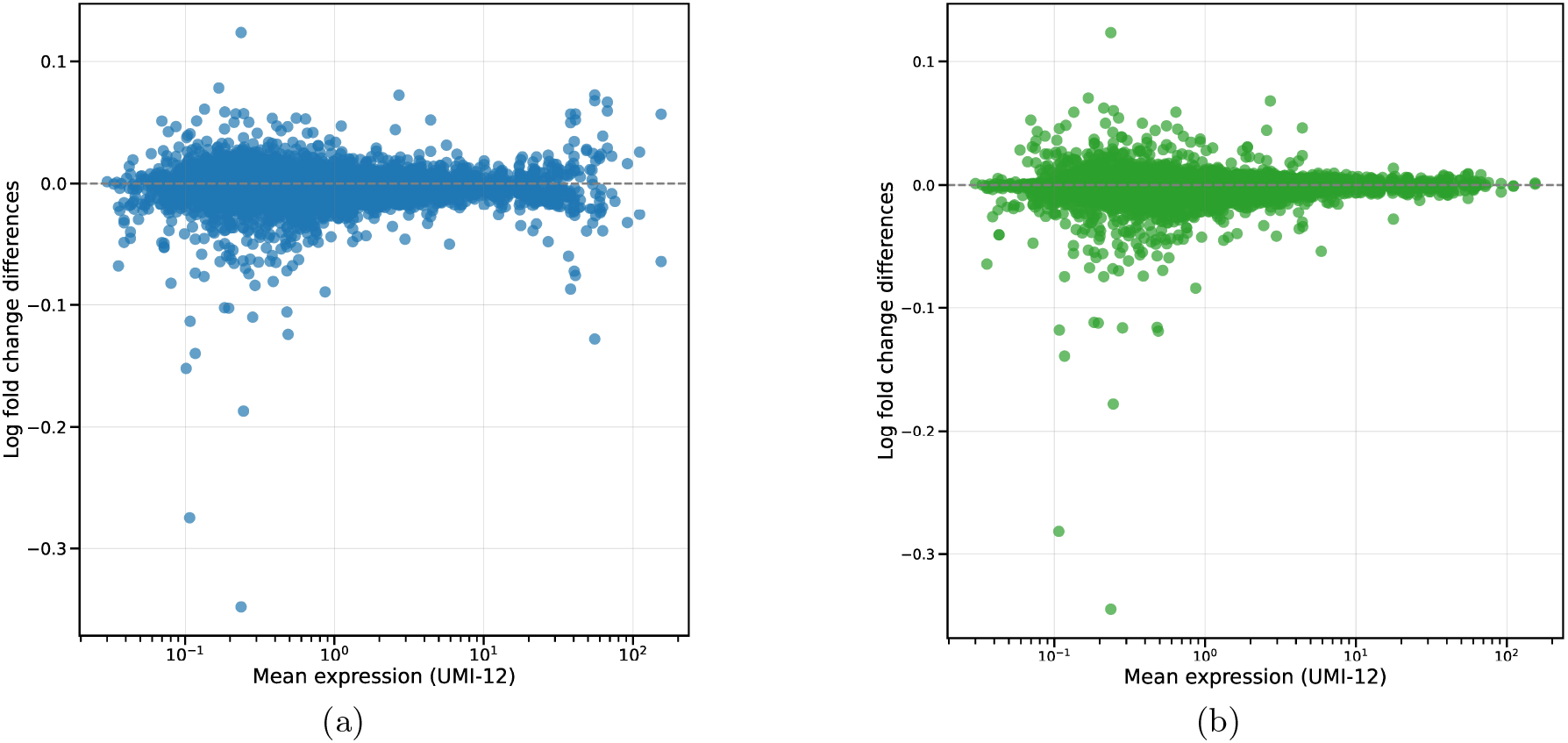
Mean expression versus LFC for all DE genes (no average expression filter) for 1k PBMC dataset. Difference between true LFC and predicted LFC using naive estimator and method-of-moments estimator for a 5-bp UMI. We filter for genes with significant p-value and moderate fold change. Performance is similar for the two methods at low expression levels, as these genes often do not even require the use of UMIs (Figure S10). As mean expression increases however, our estimator shows its improved performance (highlighted in Figure 5d,e). **a)** Log-fold changes between ground truth and naive estimator for all genes. **b)** Same as **a** but for our collision-aware estimator.

## S3 Maximum Likelihood Estimator in Poissonized Setting

Recall our statistical model: *N* identical balls are randomly assigned into *K* bins, with the observation *Y* denoting the number of bins with at least one ball. In our sequencing model, this translates to *N* mRNA transcripts before PCR amplification, *K* = 4^*k*^ possible UMIs, and *Y* unique UMIs observed. The balls (transcripts) follow some distribution *p* over the set [*K*] = {1, 2, …, *K*}, and are independent and identically distributed. Then, recall Equations (1) and (2):

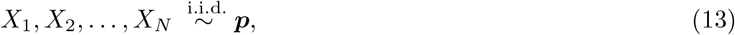

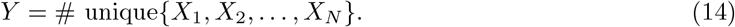

Mathematically, *Y* = |{*X*_1_, *X*_2_, …, *X*_*N*_ }|, where |*S*| denotes the cardinality of a set *S*, in our case the number of distinct UMIs observed. Given *p* and *Y*, we want to estimate *N* . By representing *Y* as a sum of *K* indicators, we can compute its mean as in Equation (3)

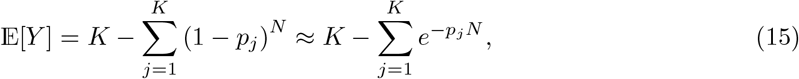

where 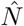 is computed as a method-of-moments estimator by inverting the above relationship.

### S3.1 Sufficient statistics

Here, we note that we do not simply observe *Y*, but in fact observe some vector *W* ∈ ℕ^*K*^, where *W*_*j*_ denotes the number of reads assigned to UMI *j* (after PCR amplification for each *X*_*i*_). However, since we do not have a good model for PCR amplification, we ignore this information and instead only consider the signature *Z* of *W*, which is

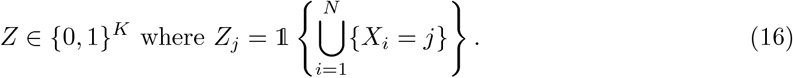

Note that 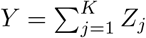. However, when *p* is not uniform, *Y* is not a sufficient statistic for *N* . The likelihood of our observations depends on the full vector *Z*, not just its sum *Y* .

#### Intuition

Consider the case where we only have 2 bins (*K* = 2), where a ball is thrown into the first one with *p*_1_ = 0.99, and the second with *p*_2_ = 0.01. Then, if we observe occupancy [0, 1], we are *reasonably* confident that this means that *N* = 1, as: ℙ(*Z* = [0, 1]|*N* = 1) = 0.01, while ℙ(*Z* = [0, 1]|*N* = 2) = .01^2^. However, if we observe *Z* = [1, 0], then our guess for *N* should be much larger, since it is likely that this consists of many balls all going into the first bin. Concretely, ℙ(*Z* = [1, 0]|*N* = 1) = 0.99, while ℙ(*Z* = [1, 0]|*N* = 10) = 0.99^10^ ≈ 0.904, i.e. the likelihood decreases much more slowly as *N* increases. Formally, writing out the likelihood shows that it does not factor into a function of *Y* and a function independent of *N*, but rather depends on the full vector *Z*.

### S3.2 Poissonized Setting

Analyzing the original model is difficult, since the indicators *Z*_*j*_ are dependent. To simplify the analysis, we consider the Poissonized model where instead of *N* balls, we have *N*^*′*^ ∼ Poisson(*N* ) balls. This is a common algorithmic analysis technique, and is a good approximation when *N* is large. Then, the number of balls in each bin *j* is independent and distributed as Poisson(*p*_*j*_*N* ), due to Poisson thinning. In this case, the indicators *Z*_*j*_ are independent Bernoulli random variables with

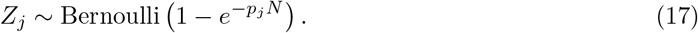

Then, the likelihood is

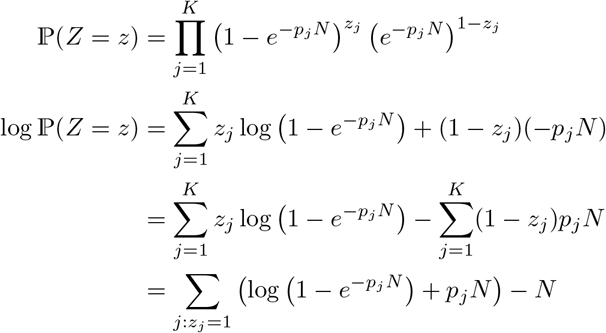

We can minimize the negative log-likelihood to obtain the MLE for *N* (gradient descent for a single parameter). Standard results show that the MLE is consistent and asymptotically normal, with asymptotic variance given by the inverse Fisher information. The Fisher information is (after checking the necessary regularity conditions):

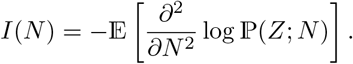

First, we compute the score function:

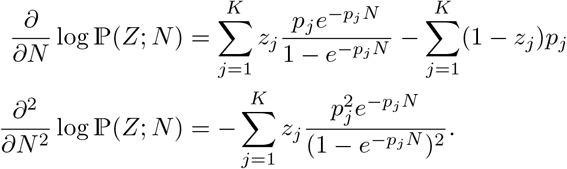

Since 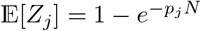, the Fisher information is

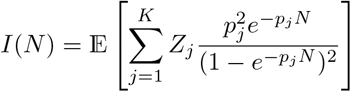

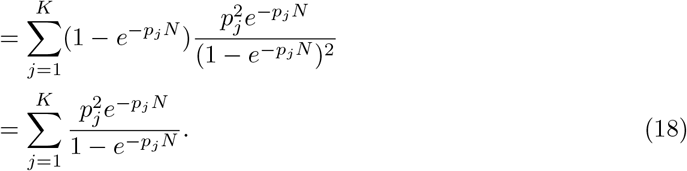

Thus, if we compute the MLE 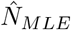 by minimizing the negative log-likelihood, we have that 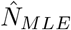 is asymptotically normal with mean *N* and variance approximately equal to the inverse of the Fisher information:

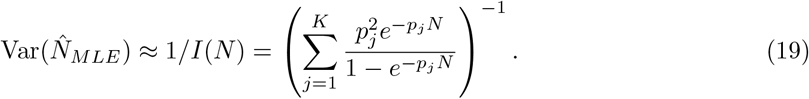

#### S3.2.1 Method of Moments Estimator in Poissonized Setting

In the Poissonized setting, the crossterm in the variance of *Y* vanishes, since the *Z*_*j*_ are independent. Thus, we have

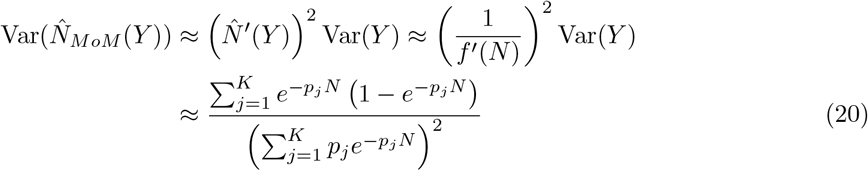

We can compare the asymptotic variances of the MLE and MoM estimators in the Poissonized setting. We show that 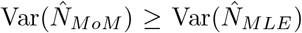, i.e., the MLE is always at least as efficient as MoM. This is equivalent to showing that

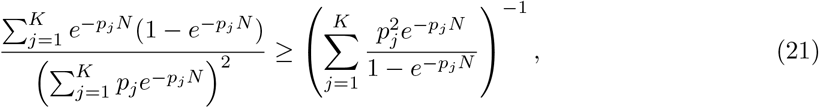

i.e., that

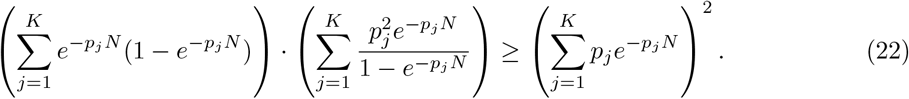

Let 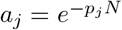 and 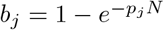. We apply the Cauchy-Schwarz inequality with

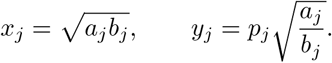

The 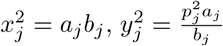, and *x*_*j*_*y*_*j*_ = *p*_*j*_*a*_*j*_. By Cauchy-Schwarz, 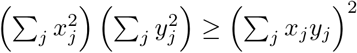 which gives exactly (22).

Equality holds when *x*_*j*_*/y*_*j*_ is constant across all *j*, i.e., when 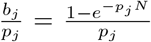 is constant. This happens if and only if *p*_*j*_ is constant across all *j*, i.e., when ***p*** is uniform, validating the observation that when ***p*** is uniform, *Y* is a sufficient statistic and the MoM estimator is efficient. When ***p*** is not uniform, the MLE outperforms the MoM estimator, with performance gap increasing as ***p*** becomes more skewed.

## S4 Empirical validation of different UMI estimates and collision-aware estimators

In this work, we have proposed degrees of estimator complexity, ranging from no UMI collision correction, to our method-of-moments estimator, to the poissonized MLE. The latter two depend on the estimated UMI distribution, which we provide models of increasing fidelity for: uniform distribution, constant PWM (cPWM), and general PWM, the latter two optionally incorporating our synthesis failure model (monotonic synthesis failure probabilities used). Additionally, the full empirical UMI distribution can be used. We discuss these two axes of improvement separately, and show that while transcript abundance estimation improves as our estimator or UMI model gets more complex, almost all the gains can be obtained by using our method-of-moments estimator with the constant PWM.

### S4.1 UMI distribution modeling

We begin by showing in Figure S12 the accuracy of different UMI distribution models, of increasing complexity, for fixed *k* = 5. We compute the L1 distance between the empirical UMI distribution (summed across all genes) and the model UMI distribution, for the 1k PBMC dataset. We analyze this for *k* = 5 since for larger *k*, many UMIs are not observed at all, and so the L1 distance is dominated by the large number of UMIs with 0 empirical probability. Next, we generate a scatter plot for each of these methods for *k* = 12 (Figure S13), where each point corresponds to a UMI, with x-coordinate given by the empirical probability of that UMI, and y-coordinate given by the probability of that UMI under the model. This shows how accounting for synthesis failure yields improved accuracy for the high probability UMIs with many trailing Ts.

**Fig. S12:**
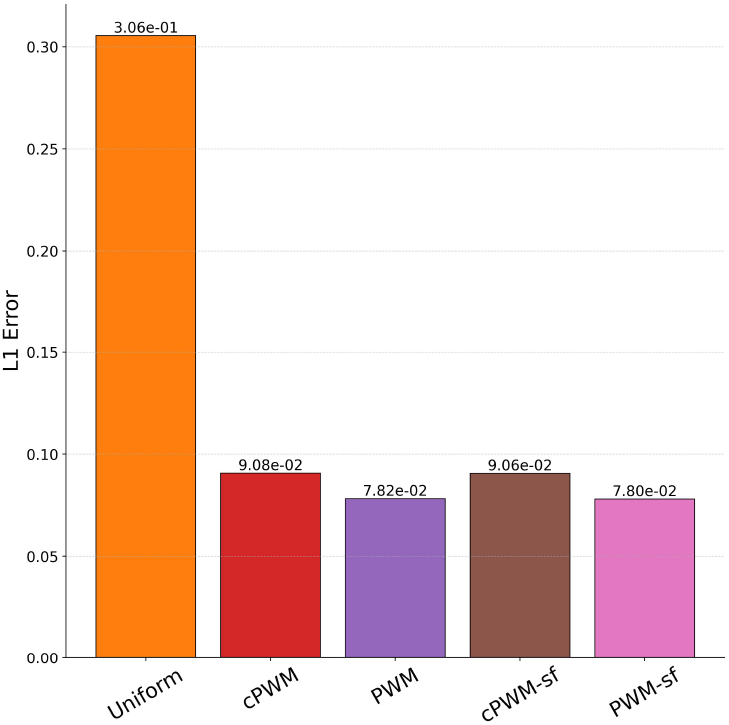
Accuracy of different UMI distribution models. We compare the accuracy of different UMI distribution models, of increasing complexity, for *k* = 5. Uniform is the simplest and performs poorly, but there are extremely diminishing returns as we increase the complexity of our UMI distribution model, with the constant PWM performing almost as well as the general PWM incorporating synthesis failure.

**Fig. S13:**
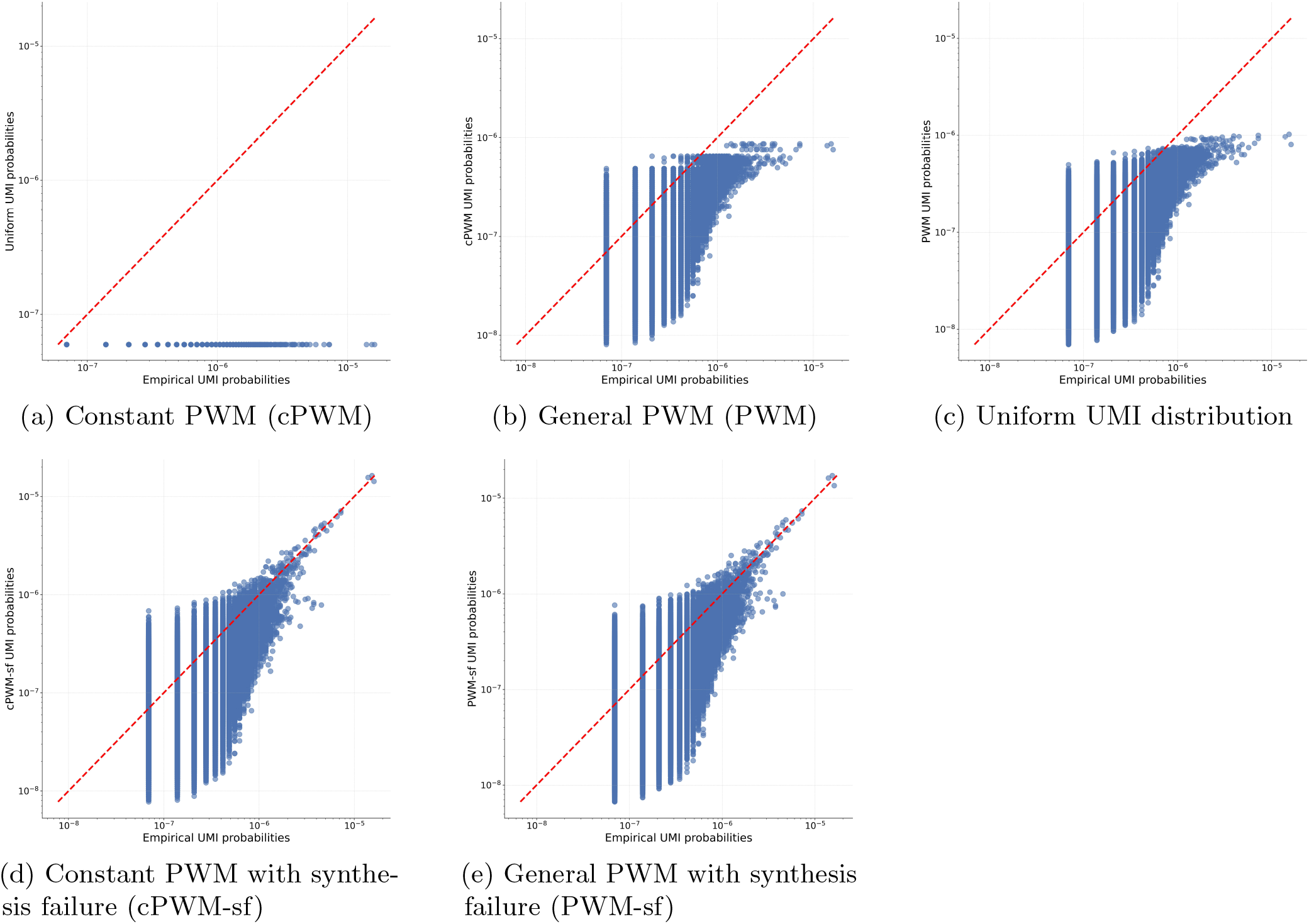
UMI modeling accuracy for PBMC 1k dataset for *k* = 12, summed across all genes. Each point corresponds to a UMI, with x-coordinate given by the empirical probability of that UMI, and y-coordinate given by the probability of that UMI under the model. The uniform model clearly performs poorly,

### S4.2 Empirical validation of poissonized MLE

We now study the interplay between the estimator and UMI distribution model complexity, by comparing the performance of the MLE and MoM estimators across different UMI distribution models (Figure S14). We see that the naive estimator performs very poorly for *k* = 5, with dramatic improvements afforded by the MoM estimator. However, the MLE only provides a very minor improvement over the MoM estimator, and almost all of the gains can be obtained by using the MoM estimator with the constant PWM. We show the performance per cell, gene, in Figure S15.

**Fig. S14:**
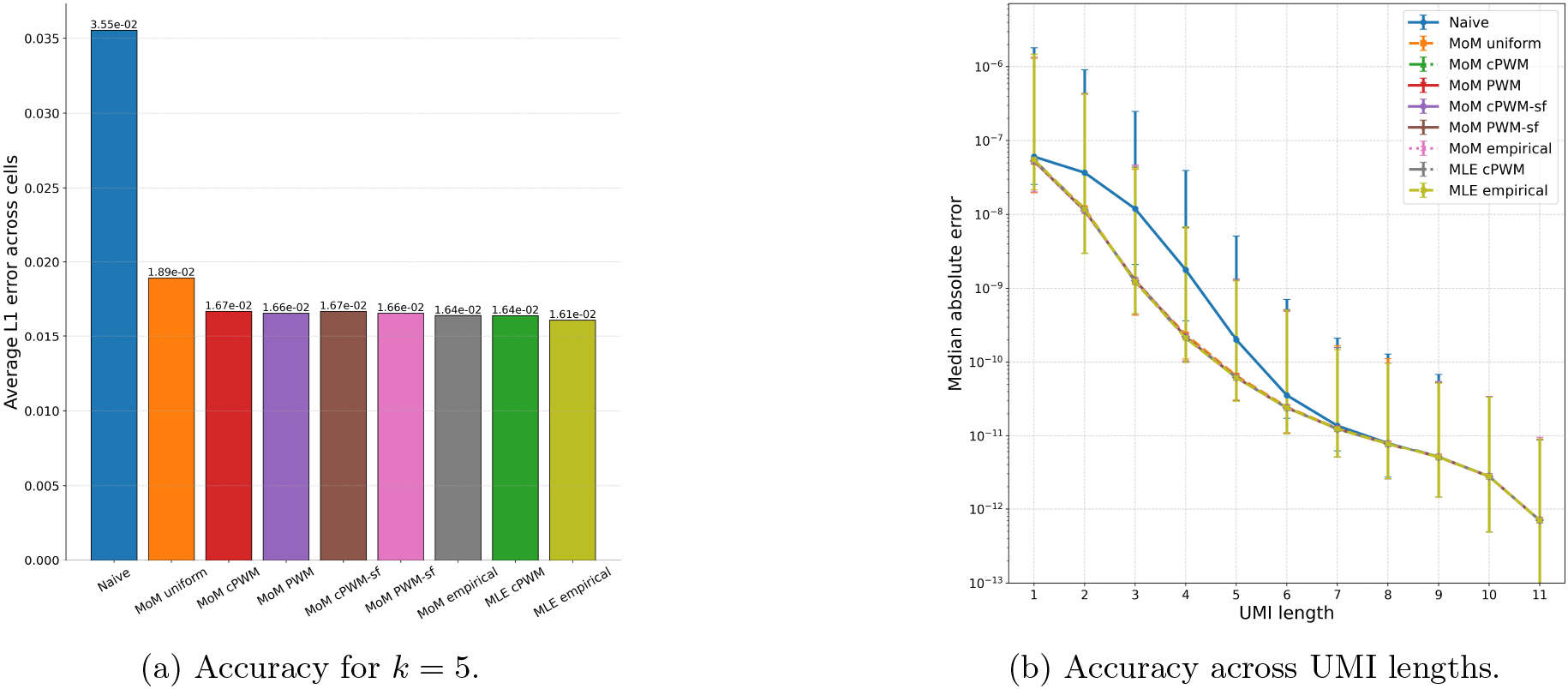
Accuracy of different estimators (naive, method-of-moments, and MLE) across different UMI distribution models.

**Fig. S15:**
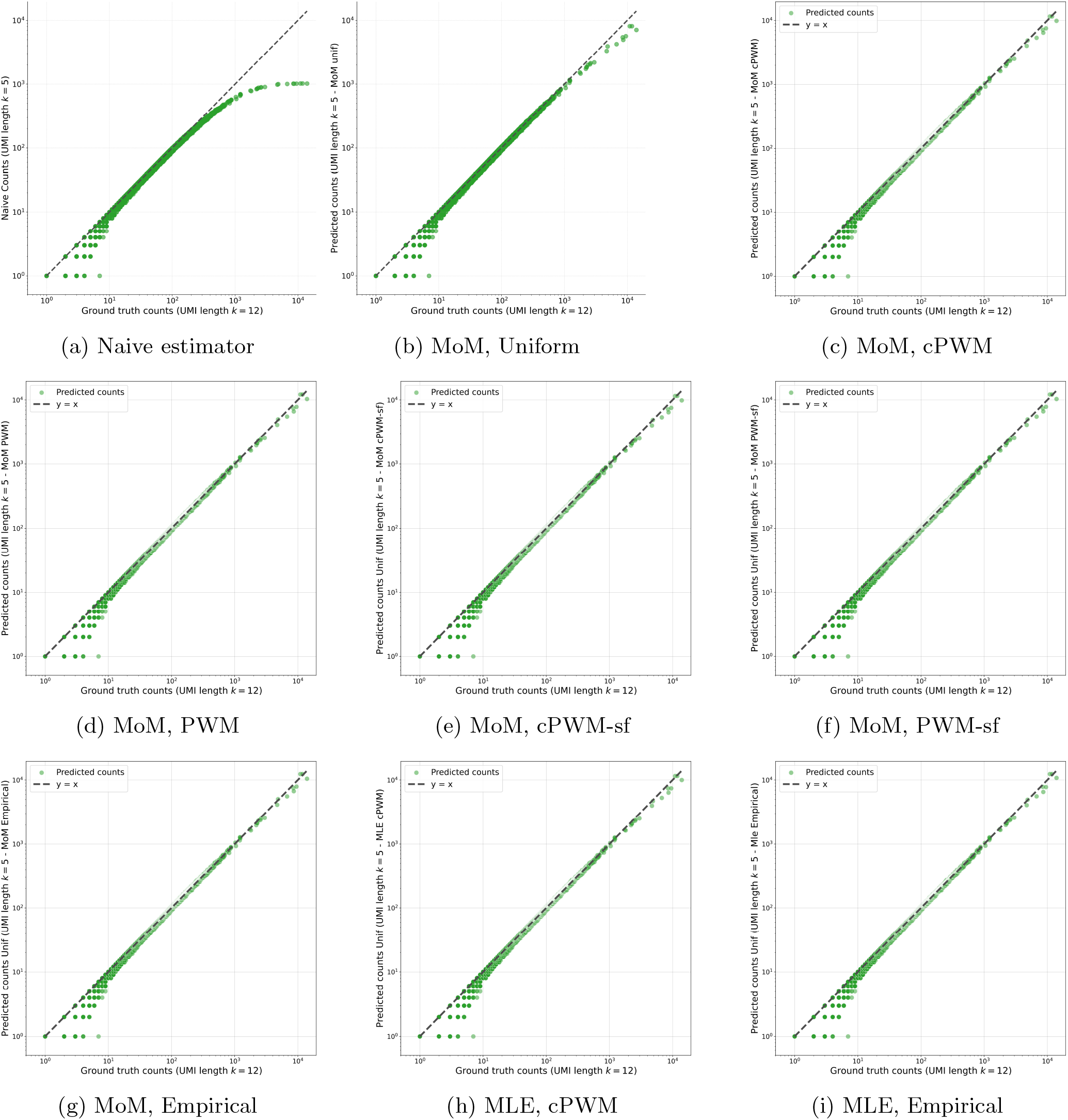
Each panel shows predicted vs. ground-truth counts for a specific estimator × UMI model combination at *k* = 5. Dramatic gains by accounting for collisions, then by using a nonuniform UMI distribution. Minimal gains from using the MLE over the MoM estimator, and from using more complex UMI distribution models.

## S5 Theoretical analysis for the method-of-moments estimator

In this section we provide additional theoretical results regarding our method-of-moments estimator, and give all deferred proofs.

### S5.1 Estimator convexity

#### Proposition 1.

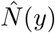 *is convex*.

*Proof* For *Y < K*, 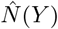 is a convex function of *Y*, as it is − log(·) composed with an affine function. Examining the edge case of *Y* = *K*, we show that the derivative is strictly increasing, in that:

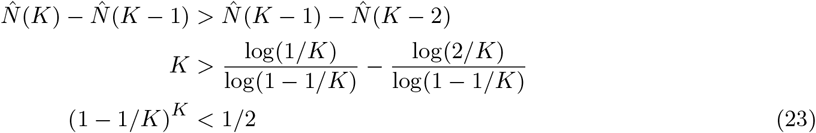

This holds true for all *K >* 1, and so the estimator is convex. This implies that defining 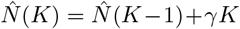 would retain the convexity of the estimator for any *γ >* ln(2).

#### S5.1.1 Extension to nonuniform UMI distributions

As before, the case of *y* = *K* is a priori undefined, as *Y*_*n*_ *< K* for all *n*. Here, we use a quadratic extrapolation, to yield a simple estimate that retains the convexity of our estimator. Quadratic extrapolation here studies finite differences, i.e. the behavior of 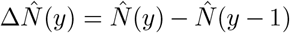. To obtain our quadratic extrapolation, we analyze the second finite difference 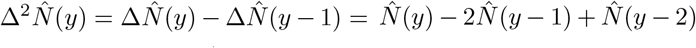. To perform quadratic extrapolation, we want 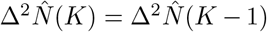. This simplifies as:

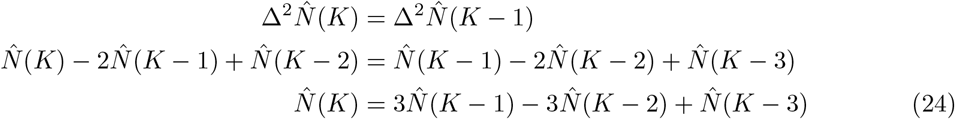

This retains the convexity of our estimator, even in the nonuniform *p* setting, as can be verified by computing that 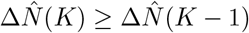.

### S5.2 Variance of MoM estimator

We compute the variance of our estimator 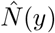 using the delta method, a first order Taylor Expansion. Let *f* (*N* ) = *Y*_*N*_ be the function relating the true number of transcripts *N* to the observed count *Y*_*N*_ = 𝔼 [*Y* ]. Recall that

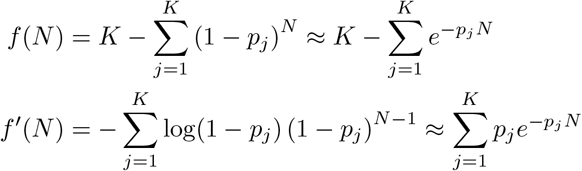

where in the last line we used that log(1 + *x*) ≈ *x* for small *x*, (1 − *x*)^*N*^ ≈ *e*^−*xN*^ for small *x*, and that *N* is large. Using the delta method, we evaluate 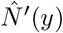, the derivative of the inverse function, evaluated at the observed count *y*. Noting that 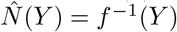 up to the linear interpolation we perform, 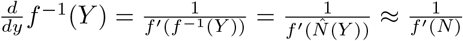, where by construction of our estimator 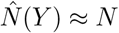.

In the uniform case, where 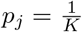 for all *j*, this simplifies as 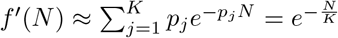. The variance of *Y* is given in Equation (4), and evaluating for general *p* yields:

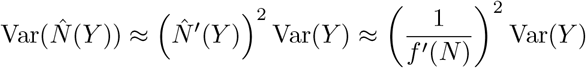

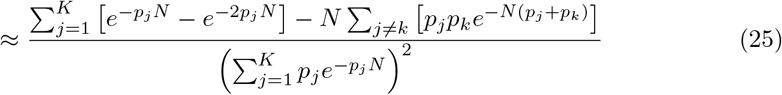

This enables us to compute confidence intervals for our predictions 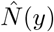, leveraging the asymptotic normality of *Y* [15] discussed in the next section.

### S5.3 Runtime and memory analysis of Method-of-Moments Estimator

We analyze the runtime and memory complexity of Algorithms 1 and 2 in terms of the total number of UMIs *K* and the maximum observed count *Y*_max_. Let 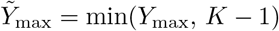.

#### Algorithm 2 (Precomputation)

The repeat loop (lines 3–6) iterates at most *n*_max_ times, where

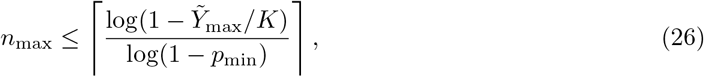

and *p*_min_ = min_*j*_ *p*_*j*_. Each iteration updates the running powers (1 − *p*_*j*_)^*n*^ for *j* = 1, …, *K* via a single multiplication per coordinate *j* and accumulates *Y*_*n*_, requiring *O*(*K*) compute per iteration. The subsequent for loop (lines 9–13) sweeps over 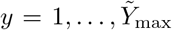 while advancing a pointer *n*_ptr_ that increases monotonically from 1 to at most *n*_max_. We have that *n* ≤ *n*_max_, and so this loop runs at most *O*(*n*_max_) times, with each interpolation step costing *O*(1). The overall time complexity of Algorithm 2 is therefore

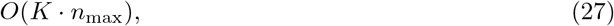

and it requires *O*(*K* + *n*_max_) memory to store the running powers (1 − *p*_*j*_)^*n*^ and the sequence 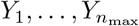. Note that this memory cost can be made independent of *K* by simply recomputing all terms at each iteration. In the uniform case *p*_*j*_ = 1*/K*, the bound simplifies to 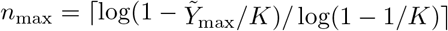.

#### Algorithm 1 (Collision-aware estimator)

Given the precomputed lookup table 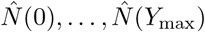, correcting each observed count requires a single *O*(1) table lookup, so the per-entry cost of applying the correction is constant. The total end-to-end complexity is

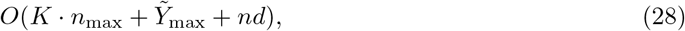

where *nd* can be replaced with the sparsity *S* by simply applying this estimator to the nonzero entries. For e.g. *k* = 6 with *K* = 4096, and 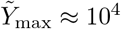 for most datasets, the precomputation cost is modest relative to the size of the counts matrix.

## S6 Asymptotic normality analysis

In this section, we leverage the asymptotic normality of *Y* to analyze the performance of our estimators in greater detail. For simplicity and concreteness, we focus on the uniform UMI setting, where *p*_*j*_ = 1*/K* for all *j* ∈ [*K*]. All results extend naturally to the nonuniform case (and the nonuniform method-of-moments estimator), but with more cumbersome expressions and less clear insights.

It is known that *Y* is asymptotically normal whenever Var(*Y* ) → ∞ [15, 23]. Thus, for *N, K*→ ∞ we have under this condition that:

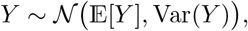

with variance (from Equation (4)), simplified for the uniform case to:

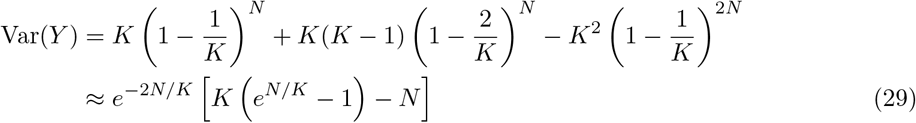

This variance vanishes for large *N* ; specifically, as we show in Proposition 2, for *N* ≥ *cK* log *K* with *c >* 1 we have Var(*Y* ) → 0 as *N, K* → ∞. In this case, *Y* converges to a point mass at *K*, and so asymptotic normality does not hold.

For *N* below this threshold, we define *f* (*N* ) = *K* 1 − *e*^−*N/K*^ ≈ 𝔼 [*Y* ]. This approximation uses log(1 − 1*/K*) ≈ −1*/K* for *K* ≫ 1, and is asymptotically tight. Then

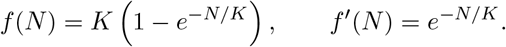

With this function-based reparameterization, we can express our (uniform UMI) estimator as 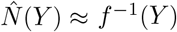, up to the log(1 − 1*/K*) and exponential approximations. This enables us to approximate the bias and variance of our estimator using the delta method.

### S6.1 Asymptotic variance of MoM estimator

Since 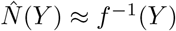, by the delta method we have

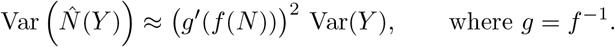

Since

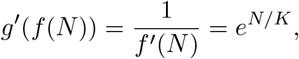

we obtain the asymptotic variance

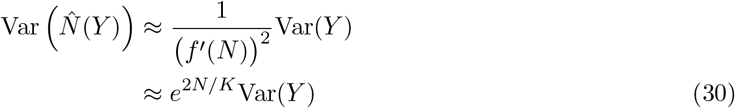

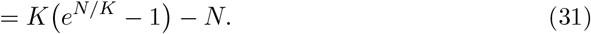

This expression governs the behavior of 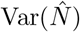 across the relevant regimes of *N* :

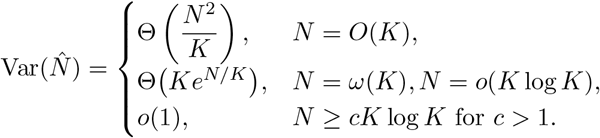

In particular, for *N* ≤ *cK* log *K* with any *c <* 1, the delta–method approximation is accurate and (31) describes the variance growth of the collision-aware estimator.

Most helpful for comparative analysis when *N* = *o*(*K* log *K*) is the characterization in Equation (30): 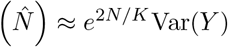. This indicates that, firstly, the variance of our estimator is always larger than the variance of *Y*, the naive estimator. However, even for large *N* approaching *K*, the variance of our estimator is only a constant factor larger than the variance of *Y* . Concretely, for *N* = *K*, we have that Var 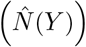 is only a factor of *e*^2^ ≈ 7.39 larger than Var (*Y* ). It is not until *N* approaches *K* log *K* that the variance of our estimator becomes significantly larger than the variance of *Y*, where in the regime of *N* = *cK* log *K* for *c <* 1, the variance of our estimator is approximately *K*^2*c*^Var (*Y* ). However, in this regime, *Y* is already highly concentrated around *K*, and so the variance of *Y* is decaying as *e*^−*N/K*^. Next, we analyze the bias of our estimator, showing that this small increase in variance enables a significant reduction in bias compared to the naive estimator.

### S6.2 Asymptotic bias of MoM estimator

We can similarly approximate the bias of our estimator 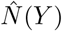 using a second-order delta method, working directly with the parameterization *f* (*N* ) = *Y*_*N*_, where *Y*_*N*_ = 𝔼 [*Y* ].

We compute the second derivative of the inverse function *g* = *f* ^−1^ using the chain rule:

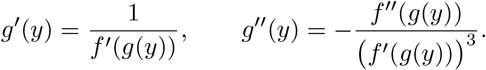

Evaluating at *y* = *f* (*N* ) (so *g*(*f* (*N* )) = *N* ) yields

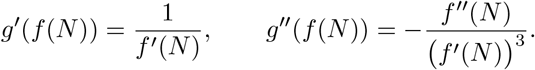

The second-order delta method gives

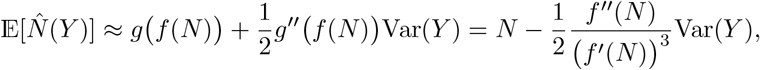

so the bias of the method-of-moments estimator is

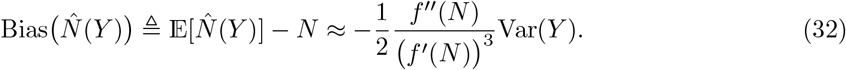

Using the approximations above,

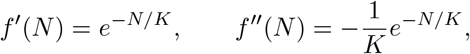

and so

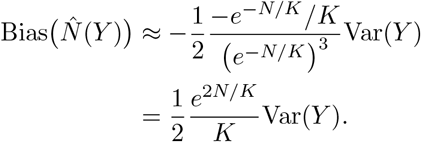

Using the variance approximation from (29), for when *N < K* log *K*,

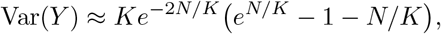

we obtain the explicit bias, noting that when *N* ≥ *K* log *K, Y* = *K* with high probability (Proposition 2), and so the bias is Θ(*N* ):

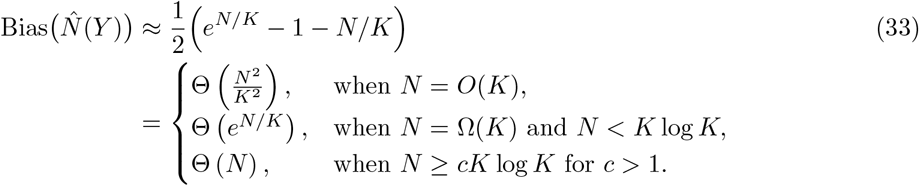

For *N* ≤ *K*, the bias is smaller than both 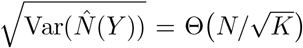 and the *O*(*N* ^2^*/K*) bias of the naive estimator. Thus, in all pre-saturation regimes, the MoM estimator is only very mildly upward biased. However, once *N > K*, the bias starts increasing exponentially with *N/K*, maxing out at Θ(*N* ) once *N* reaches *K* log *K* and *Y* saturates.

### S6.3 Extension to nonuniform UMIs

For a nonuniform UMI distribution ***p***, we can define *f* accordingly and proceed with the analysis:

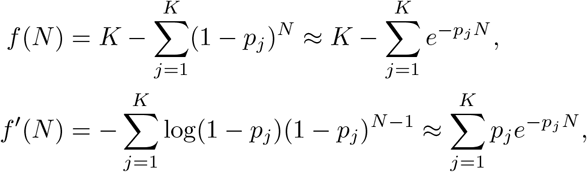

where the approximations use log(1 + *x*) ≈ *x* and (1 − *x*)^*N*^ ≈ *e*^−*xN*^ for small *x*, and that *N* is large. Differentiating once more gives

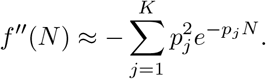

Let *g* = *f* ^−1^ denote the inverse function. Then 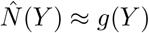, and *Y* is approximately Gaussian with mean *f* (*N* ) and variance Var(*Y* ). Using the approximations above,

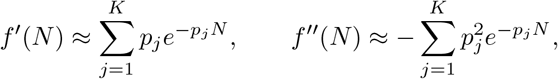

and (32) becomes

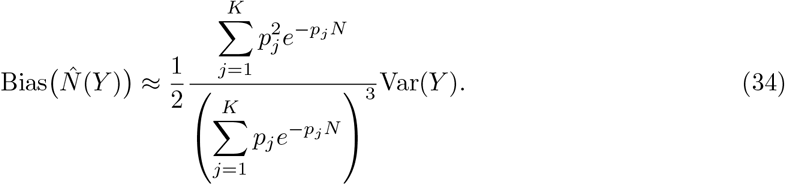

Similarly,

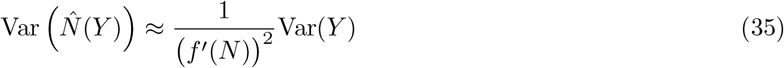

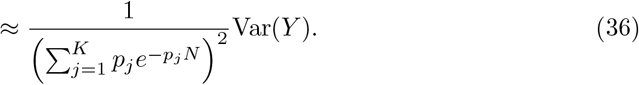

## S7 Optimality of method-of-moments estimator for uniform UMIs

On its surface, our proposed estimator seems quite simplistic. It only matches the first moment of *N*, and fails to take into account any higher order moments of *Y* . Additionally, since the estimator is a convex function of *y*, by Jensen’s inequality 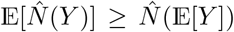, implying that 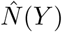 will overestimate *N* . However, as we show, this estimator yields good estimation up until the threshold of impossibility.

### S7.1 MSE analysis and comparison

One interesting note is the directions of the biases. The naive estimator is always negatively biased, as collisions cause undercounting. On the other hand, our estimator is positively biased, as it is a convex function of *Y*, where our moment matching condition combined with Jensen’s inequality imply that 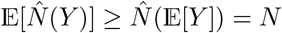.

Recall that the variance of *Y* is

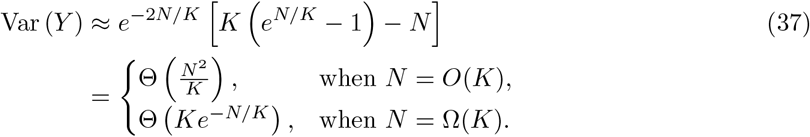

The bias of the naive estimator is:

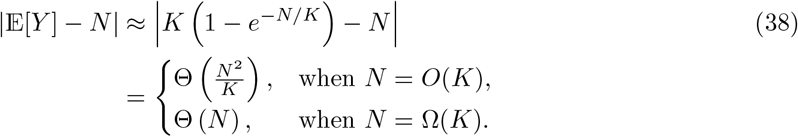

The MSE of the naive estimator is always dominated by the squared bias term. Splitting our analysis into regimes (tabulated in Table 1), we have that:

1. 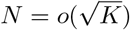. Here, both estimators have vanishing bias and variance.
2. *N* = *o*(*K*), 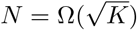 .Here, the Θ(*N* ^2^*/K*) bias of the naive estimator starts growing, leading to an MSE dominated by the squared bias of order Θ(*N* ^4^*/K*^2^). Our MoM estimator still has vanishing bias, but Var (*Y* ) ≈ *O*(*N* ^2^*/K*). This retains MSE sublinear in *N* .
3. *N* = Ω(*K*), *N* = *o*(*K* log *K*). The naive estimator already has linear bias, and so an MSE of order *N* ^2^. Here, V ar (*Y* ) ≈ *Ke*^−*N/K*^ is decaying, but our estimator now has inflated variance, leading to Var 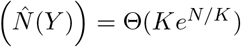, dominating the MSE of our MoM estimator.
4. *N* ≥ *cK* log *K* for *c >* 1. Here, *Y* has saturated (by Proposition 2), and so both estimators have bias and MSE on the order of *N* ^2^.

**Table S1:**
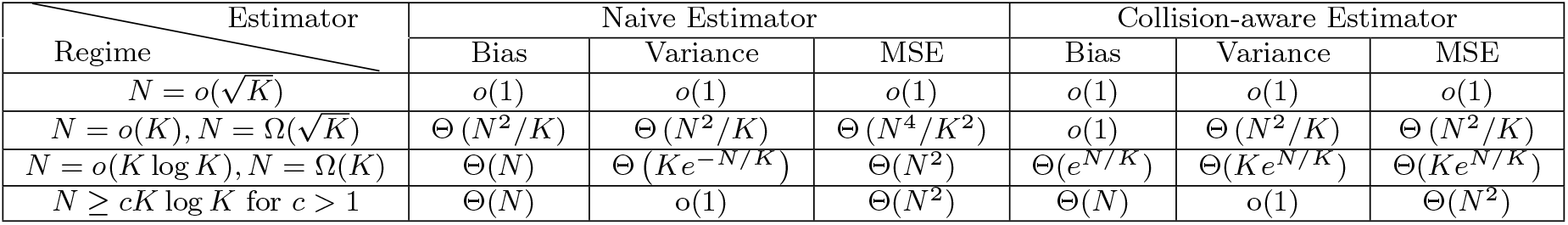
Comparison of Bias, Variance, and MSE for naive and collision-aware estimators across different regimes of *N*. Table replicated from Table 1 in the main text for convenience.

This highlights that when 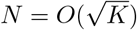 both estimators attain vanishing MSE. However, for larger *N*, the naive estimator has increasing bias which dominates the MSE. When *N* falls between *K* and *K* log *K*, the MSE of the naive estimator already scales as *N* ^2^, dominated by the bias, while our estimator performs slightly worse than the earlier MSE of Θ(*N* ), but still sub-quadratic. The high level takeaway is that in order to minimize the MSE, *K* is currently taken so that *N* ≤ *K*^2*/*3^ to avoid the large bias of the naive estimator. However, *K* can in fact be selected such that *N* is a constant multiple of *K*, in which case the MSE is still *O*(*N* ). Next, we discuss impossibility results and lower bounds in this setting.

### S7.2 Impossibility beyond *N > K* log *K*: proof of Proposition 2

From the classical coupon collector problem, it is known that the expected number of balls required until all bins are filled is *N* = *K* log *K* + *O*(*K*). This threshold is tight: taking *N* to be larger than *K* log *K* by any multiplicative constant yields vanishing (with *K*) probability of observing *Y < K*.

#### Proposition 2

*For N* = *cK* log *K with c >* 1, ℙ (*Y* = *K*) ≥ 1 − *K*^1−*c*^.

*Proof of Proposition 2* Define the indicator random variables *Z*_*j*_ = 𝟙{∪_*i*_{*X*_*i*_ = *j*}}, whether bin *j* is filled, for *j* ∈ [*K*]. Then, *Y* = ∑_*j*_ *Z*_*j*_ .

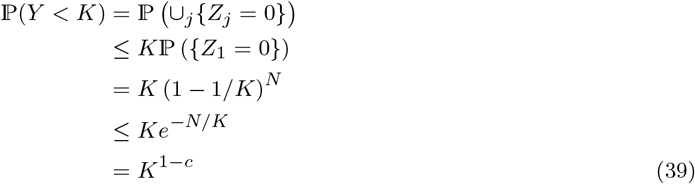

where a union bound is used, followed by the inequality that 1 − *x* ≤ *e*^−*x*^.

Since *Y* will be equal to *K* with high probability, we cannot distinguish between *N* = 2*K* log *K* and *N* ^*′*^ = *K*^3^, for example, and so *N* cannot be estimated to any nontrivial accuracy. Extending this to the nonuniform *p* setting is difficult, without a closed form solution. Defining *p*_min_ = min_*j*_ *p*_*j*_, we see that *N* = Ω(1*/p*_min_) is necessary, as otherwise the UMI corresponding to *p*_min_ will not have been observed with high probability. Conversely, *N* = *O*(log(*K*)*/p*_min_) is sufficient, by a similar union bounding argument. Again, by concavity, *N* = Ω(*K* log *K*) is necessary.

#### S7.2.1 Extending saturation threshold beyond *K* log *K* by adjusting the UMI distribution

The above observations regarding the saturation threshold scaling as 1*/p*_min_ imply that we can increase our threshold for feasible estimation by decreasing the minimum probability. A natural question is then: how far can we extend our nontrivial estimation? As we drop *p*_min_ → 0, *Y* will not saturate until 1*/p*_min_, but gain increased variance before. Theoretically, given a known distribution of true UMI counts (*N* ), or a range of feasible *N*, one could compute the best UMI distribution with respect to this Bayes Risk or Minimax Risk. Even by taking a simple setting, where per-nucleotide probabilities are [1 − *x*, 1 − *x*, 1 − *x*, 3*x*] for [A,C,G,T], with *x* ∈ [0, 1*/*3], by taking *x* to 0 we can guarantee that *Y* will not saturate, and that via the asymptotic normality argument below, we can compute the MSE for any fixed *x* and *N* .

### S7.3 Cramér–Rao lower bound in binomial setting

To begin, *N* is a discrete parameter, and so we technically cannot directly apply the Cramér–Rao lower bound. However, since the likelihood is a continuous function of *N*, we can instead perform inference when *N* ∈ R_+_ and then apply the Cramér–Rao lower bound to the continuous function.

Recall the indicator random variable based definition of *Y*, with *Z*_*j*_ = 𝟙{∪_*i*_{*X*_*i*_ = *j*}}, whether bin *j* is filled, for *j* ∈ [*K*], and *Y* = ∑_*j*_ *Z*_*j*_. Here,

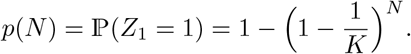

In our setting these *Z*_*j*_ are correlated, making the analysis difficult, so we approximate *Y* by 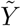, a binomial where the *Z*_*j*_ are independent:

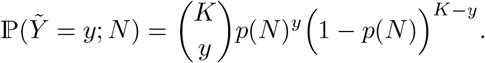

The mean of 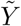 matches *Y*, as:

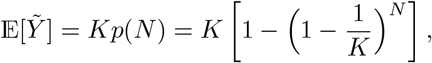

and its variance is (in this binomial approximation)

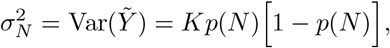

which is approximately the variance of *Y* .

The log-likelihood for 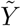 is then

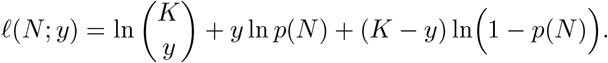

The Fisher information is defined as

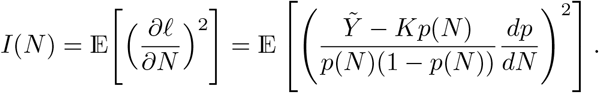

Plugging in for the variance of 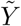 in the numerator, it follows that

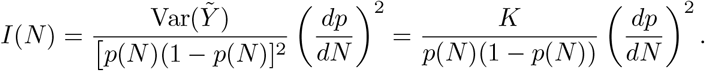

For large *K*, we may use the approximations

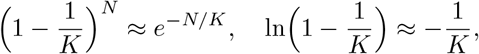

so that

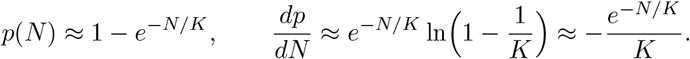

Then, the Fisher information becomes

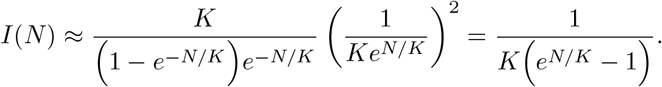

This allows us to state our Cramér–Rao lower bound:

#### Theorem 1.

*Any unbiased estimator* 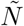 *of N, given observation* 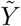, *satisfies*

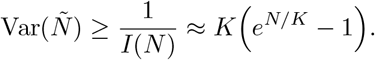

Our method-of-moments estimator matches this lower bound in this simplified setting (Equation (31)), highlighting the near-optimality of our estimator. Observe that our estimator has nontrivial bias for *N* = Ω(*K*), and so this lower bound does not directly apply in this regime, but for *N* = *o*(*K*) our estimator is asymptotically unbiased. Evaluating this expression, we have that as *N* approaches *K* ln *K*, we have *e*^*N/K*^ ≈ *K*, so that 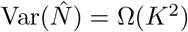, indicating a dramatic increase in estimation error. In this regime, where nearly all bins are occupied, even small differences in *Y* lead to large differences in 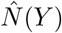, and no estimator can achieve significantly lower variance than the approximate lower bound. Our method-of-moments estimator achieves this approximate lower bound up to a constant factor, and so is near-optimal in this regime.

## Notes

### Competing Interest Statement

The authors have declared no competing interest.

### Summary of Updates

Version to be presented at RECOMB-seq 2026. Major changes include extended MALAT1 analysis, quantification and estimation of truncated synthesis, MLE estimator in poissonized setting, asymptotic normality of method of moments estimator, and additional empirical validation

